# SRSF6 balances mitochondrial-driven innate immune outcomes through alternative splicing of BAX

**DOI:** 10.1101/2022.07.18.500495

**Authors:** Allison R. Wagner, Chi G. Weindel, Kelsi O. West, Haley M. Scott, Robert O. Watson, Kristin L. Patrick

**Author notes:** Lead contact, Twitter: @The_PW_Lab (Kristin L. Patrick).

## Abstract

To mount a protective response to infection while preventing hyperinflammation, gene expression in innate immune cells must be tightly regulated. Despite the importance of pre-mRNA splicing in shaping the proteome, its role in balancing immune outcomes remains understudied. Transcriptomic analysis of murine macrophage cell lines identified Serine/Arginine Rich Splicing factor 6 (SRSF6) as a gatekeeper of mitochondrial homeostasis. SRSF6 orchestrates this by directing alternative splicing of the mitochondrial pore-forming protein BAX. Loss of SRSF6 promotes accumulation of BAX-κ, a variant that sensitizes macrophages to undergo cell death and triggers upregulation of interferon stimulated genes through cGAS sensing of cytosolic mitochondrial DNA. Upon pathogen sensing, macrophages regulate SRSF6 expression to control the liberation of immunogenic mtDNA and adjust the threshold for entry into programmed cell death. This work defines BAX alternative splicing by SRSF6 as a critical node not only in mitochondrial homeostasis, but also in the macrophage’s response to pathogens.

## INTRODUCTION

When innate immune cells like macrophages sense pathogen or damage associated molecular patterns (PAMPs or DAMPs), they rapidly induce transcription of hundreds of genes encoding cytokines, chemokines, and antimicrobial mediators (Hagai et al., 2018; Ramsey et al., 2008). While these transcripts are being synthesized by RNA polymerase II, they are subject to several critical post-transcriptional processing steps including 5’ capping, cleavage and polyadenylation, and pre-mRNA splicing, whereby introns are removed and exons are ligated together to generate mature RNAs (Carpenter et al., 2014). Pre-mRNA splicing plays a key role in global regulation of the transcriptome and thus the proteome, with 92-94% of the human genome subject to alternative splicing (Wang et al., 2008) and >80% of alternative splicing predicted to impact protein functionality (Yura et al., 2006).

Splicing regulatory proteins play a critical role in maintaining the fidelity of splicing while permitting the flexibility needed for alternative exon usage. One major family of splicing regulators is the Serine/arginine rich, or SR proteins. These proteins recognize and bind to exonic splicing enhancer sequences to define exon locations, thus directing the U snRNPs to cis-splicing signals in particular nearby introns. The SRs also function at other steps of the RNA life cycle including mRNA export, localization, decay, and translation (Howard and Sanford, 2015). Many connections have been made between SR proteins and cancer, with aberrant expression of SR proteins commonly observed in patients with multiple myeloma and acute myeloid leukemia (Liu et al., 2022; Song et al., 2019; Wan et al., 2019). There are also emerging roles for SR proteins in regulating immune homeostasis. SRSF1, the best studied SR protein, limits autoimmunity via a role in maintaining healthy regulatory T cells (Katsuyama and Moulton, 2021). SRSF3 has been shown to negatively regulate IL-1β release during *E. coli* infection of THP-1 monocytes and SRSF2 promotes herpes simplex virus replication by binding to viral promoters and controlling splicing of viral transcripts (Wang et al., 2016).

To better capture the phenotypes associated with SRSF proteins in innate immunity, we carried out a transcriptomics study of a panel of SRSF knockdown (KD) RAW 264.7 macrophage cell lines. This analysis revealed a remarkable degree of diversity in how individual SR proteins influence innate immune responses (Wagner et al., 2021). One particularly striking phenotype we uncovered was that of *Srsf6* KD macrophages, which express high basal levels of *Ifnb1* and interferon stimulated genes (ISGs). SRSF6 is a 55kDa protein that is essential for viability of *Drosophilia melanogaster* and *Mus musculus* (Mason et al., 2020; Ring and Lis, 1994)). Multiple studies report that SRSF6 preferentially binds purine-rich exonic splicing enhancers, with a predicted consensus site in humans of USCGKM (where S represents G or C; K represents U or G; M represents A or C) (Liu et al., 1998). SRSF6 activity is at least in part controlled via phosphorylation by the dual specificity kinases CLK1 and DYRK1a (Hara et al., 2013; Yin et al., 2012), which activates SRSF6 shuttling between the cytoplasm and the nucleus (Sapra et al., 2009). SRSF6 abundance has been repeatedly associated with cancer, liver disease, and diabetes (Jensen et al., 2014; Juan-Mateu et al., 2018; Li et al., 2021) and recent work has identified several roles for SRSF6 in mitochondrial function and cell death. For example, loss of SRSF6 decreases mitochondrial respiration, leading to increased cellular apoptosis in human endoC-BH1 endothelial cells, likely via alternative splicing of cell death factors like *Bim*, *Bax*, *Diablo*, and *Bclaf1* (Juan-Mateu et al., 2018). Likewise, phosphorylation of SRSF6 induces alternative splicing of mitochondria related genes (e.g. *Polg2*, *Nudt13*, *Guf1*, *RnaseI*, *Nme4*) in a mouse model of fatty liver disease as well as in human hepatitis patients (Li et al., 2021).

Here, we report that SRSF6 works to limit basal type I interferon (IFN) expression and apoptosis in murine macrophage cell lines and primary macrophages. The mechanisms underlying these phenotypes converge on alternative splicing of the pro-apoptotic factor BAX; specifically, upregulation of a BAX variant called BAX-kappa (Bax-κ). Our findings support a model whereby Bax-κ expression renders BAX mitochondrial pores permissive to mtDNA release and sensitive to triggers of programmed cell death. These studies provide insight into how BAX alternative splicing controls mitochondrial homeostasis and illuminate an unappreciated role for SRSF6 in balancing macrophage innate immune responses.

## RESULTS

### SRSF6 knockdown activates type I interferon gene expression in macrophages

To appreciate the distinct contribution of individual SR proteins to macrophage gene expression, we generated RAW 264.7 macrophage cell lines (RAW MΦ) stably expressing shRNA hairpins directed against SRSF1, 2, 6, 7, and 9, as previously reported in (Wagner et al., 2021). Most SR proteins are ubiquitously expressed across cell types, and we confirmed expression of each of these SRs in our RAW MΦ (Fig. S1A). Because it is well established that several members of the SR protein family are essential genes across multiple cell types (Feng et al., 2009; Goldberger et al., 2021; Ortiz-Sanchez et al., 2019; Wang et al., 2001; Xu et al., 2005) and because high-quality transcriptomics analyses have been carried out successfully for SR family members using shRNA or siRNA gene silencing (Consortium, 2004; Van Nostrand et al., 2020), we opted to stably knockdown these factors, instead of relying on CRISPR-mediated gene editing. To identify major transcriptomic changes due to loss of SR proteins, we isolated total RNA from each of these KD cell lines and a scramble (SCR) control that was selected alongside the SR KDs, performed RNA-seq, and measured differential gene expression using the CLC Genomics Workbench, as in (Wagner et al., 2021). Using a simple hierarchical clustering algorithm, we visualized the gene expression profiles of each SR KD macrophage cell line and pinpointed SRSF6 as unique amongst the SR proteins queried (correlation between SRSF6’s node and the rest of the tree = 0.138) (Fig 1A). Manual annotation of genes in clusters that were uniquely impacted by loss of SRSF6 revealed several downregulated genes related to mitochondrial biology (Fig. 1B). On the other hand, many upregulated genes fell into the category of interferon stimulated genes (ISGs) (Fig. 1C), as defined by (Kim et al., 2018; Liu et al., 2012; Thomas et al., 2006; Zahoor et al., 2014). ISGs are a group of genes whose transcription is activated through IFNAR receptor-mediated type I interferon signaling. Generally, very few ISG transcripts accumulate in resting macrophages. Direct visualization of RNA-seq reads using the Integrated Genome Viewer shows elevated reads across all coding exons for representative ISGs *Rsad2* (Fig. 1D) and *Mx1* (Fig. S1B). Ingenuity Pathway Analysis confirmed overrepresentation of differentially expressed genes in functional categories related to interferon and antiviral responses (Fig. 1E), indicating an overall increase in type I IFN signaling. Furthermore, RT-qPCR confirmed basal ISG expression in two independently derived *Srsf6* KD cell lines (Fig. 1H-J), with the degree of ISG accumulation correlating with SRSF6 KD efficiency and protein expression (Fig 1F-G). We also measured ISG accumulation upon transient siRNA KD of SRSF6 in RAW MΦ cells (Fig. S1C), arguing against off-target effects resulting from stable selection of KD cell lines. Importantly, this phenotype was recapitulated by *Srsf6* siRNA KD in primary cell types including bone marrow derived macrophages (BMDMs) (Fig. 1K) and mouse embryonic fibroblasts (MEFs) (Fig. S1D). Differential expression of genes involved in type I IFN responses was not observed in other resting SRSF KD RAW MΦ cell lines, suggesting this phenotype is unique to loss of SRSF6 and is not a general consequence of interfering with splicing (Wagner et al., 2021).

**Figure 1:**
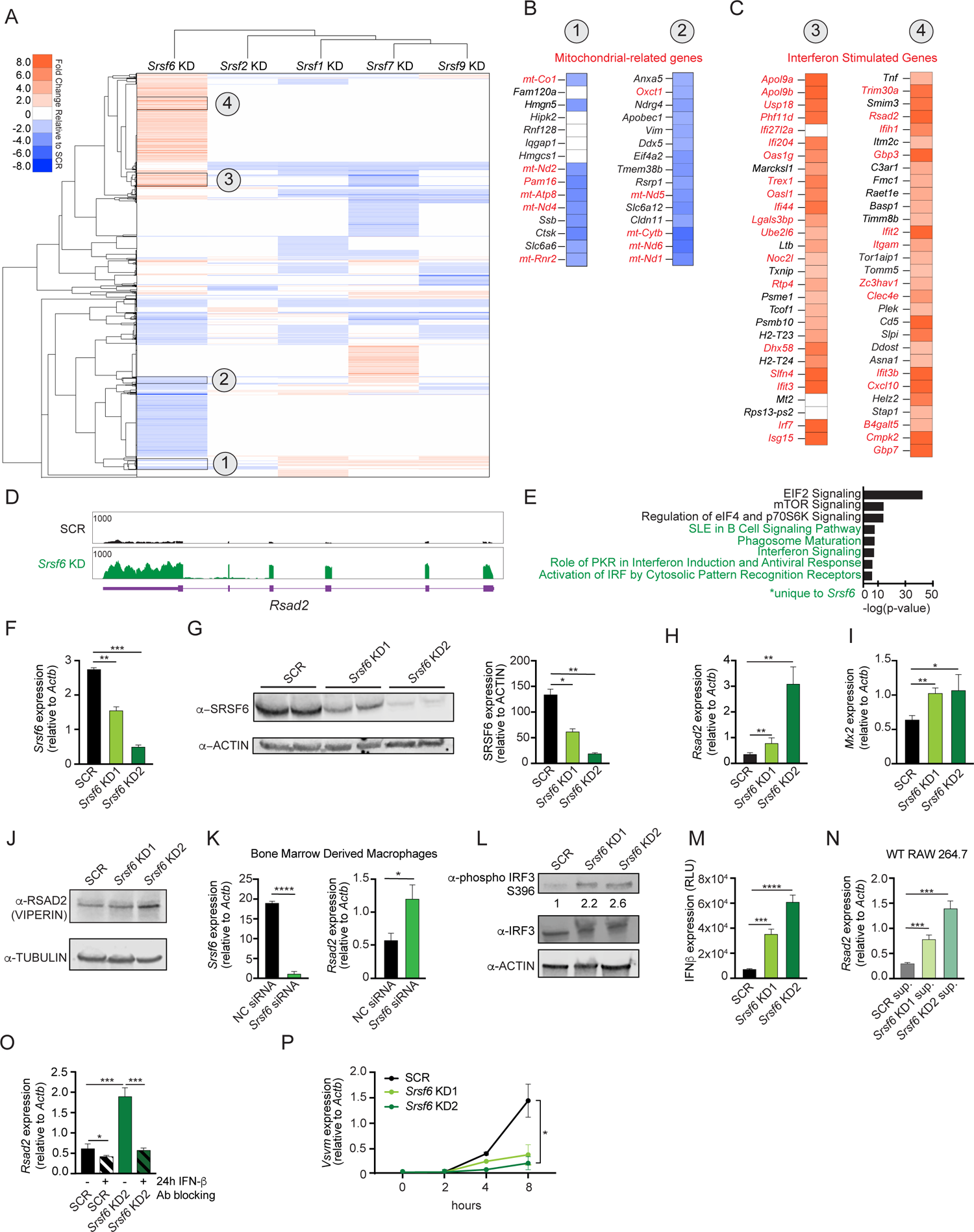
SRSF6 controls basal type I interferon expression in macrophages. **A.** Heatmap of differentially expressed genes after knockdown of SRSF1, 2, 6, 7 and 9 in RAW 264.7 macrophage-like cell lines relative to a scramble (SCR) shRNA control. **B.** Differential gene expression of mitochondria related genes (red) in *Srsf6* KD RAW 264.7 cells. **C.** As in B but highlighting ISGs (red). **D.** Integrative Genomics Viewer (IGV) tracks of *Rsad2* from *Srsf6* KD macrophage RNA seq. **E.** Ingenuity Pathway Analysis showing canonical pathways from *Srsf6* KD RAW 264.7 macrophage RNA seq. Green indicates pathways unique to SRSF6. **F.** RT-qPCR of *Srsf6* in *Srsf6* KD RAW 264.7 cells. **G.** Immunoblot of SRSF6 in *Srsf6* KD RAW 264.7 cells. **H.** RT-qPCR of *Rsad2* in *Srsf6* KD RAW 264.7 cells. **I.** RT-qPCR of *Mx2* in *Srsf6* KD RAW 264.7 cells. **J.** Immunoblot of RSAD2 (VIPERIN) in *Srsf6* KD RAW 264.7 cells. **K.** RT-qPCR of *Srsf6* and *Rsad2* in *Srsf6* siRNA KD BMDMs compared to a negative control (NC) siRNA control. **L.** As in G but for phosphorylated IRF3 and total IRF3. Numbers indicate densiometric measurements of pIRF3 (LICOR). **M.** Protein quantification of extracellular IFNβ in *Srsf6* KD RAW 264.7 cells measured by relative light units (RLU). **N.** RT-qPCR of *Rsad2* in WT RAW 264.7 cells incubated with SCR or *Srsf6* KD RAW 264.7 cells supernatants for 24 h. **O.** RT-qPCR of *Rsad2* in *Srsf6* KD RAW 264.7 cells given IFNβ neutralizing antibody treatment for 24 h. **P.** VSV replication in *Srsf6* KD RAW 264.7 cells at 0, 2, 4, 8 h post infection (MOI=1) measured by RT-qPCR of *Vsvm*. All data is compared to a scramble control unless indicated. Data are expressed as a mean of three or more biological replicates with error bars depicting SEM. Statistical significance was determined using two tailed unpaired student’s t test. *=p<0.05, **=p<0.01, ***=p<0.001, ****=p<0.0001.

Having concluded that SRSF6 plays a role in regulating basal ISG expression, we sought to investigate the cell-intrinsic vs. cell-extrinsic nature of this phenotype. Phosphorylation of IFN regulatory factor 3 (IRF3) is a critical step in the initial stages of PAMP and DAMP sensing that lead to production of IFN-β. Immunoblot analysis revealed increased levels of phospho-IRF3 (S396) in resting *Srsf6* KD macrophages (Fig. 1L). Higher levels of IFN-β protein were also measured in the supernatants of *Srsf6* KD RAW MΦ cells via ISRE reporter cells (Hoffpauir et al., 2020)(Fig. 1M). Consistent with higher levels of IFN-β secretion, supernatants from *Srsf6* KD macrophages were sufficient to stimulate ISG expression in naïve wild-type RAW MΦ cells (24 h incubation) (Fig. 1N). Elevated basal ISG expression in *Srsf6* KD cells was rescued by treatment with an IFN-β neutralizing antibody (Fig. 1O). Together, these data suggest that IRF3-mediated expression of IFN-β drives upregulation of basal ISG expression in *Srsf6* KD RAW MΦ cells. Finally, to demonstrate the biological relevance of increased SRSF6-dependent basal ISG expression, we infected cells with the single stranded RNA virus VSV, which is hypersensitive to even low levels of ISG expression (Wagner et al., 2021; West et al., 2019). We observed a dramatic restriction of VSV replication in *Srsf6* KD RAW MΦ cells compared to SCR controls (Fig. 1P and S1E), supporting a *bone fide* role for SRSF6 as a negative regulator of antiviral immunity.

### Cytosolic mtDNA triggers cGAS-dependent DNA sensing in *Srsf6 KD* macrophages

We next sought to identify the trigger of the higher basal type I IFN expression in *Srsf6* KD RAW MΦ cells. Mitochonrial DNA (mtDNA) can activate cytosolic DNA sensing pathways when mitochondria are damaged or depolarized such that mtDNA is released from the inner matrix (West et al., 2015). Because mitochondrial genes were downregulated in *Srsf6* KD RAW MΦ, we hypothesized that the mitochondrial network might be impacted by a loss of SRSF6. To begin to establish mtDNA as a source of the increased ISG expression, we first set out to determine if cytosolic mtDNA was increased in *Srsf6 KD* macrophages. To measure cytosolic mtDNA we utilized differential centrifugation to separate cytosolic and cellular membrane fractions (the latter of which include mitochondria) from SCR and *Srsf6* KD RAW MΦ (Fig. 2A). We detected significant enrichment of several mtDNA genes (*Cytb*, *Dloop2*, and *Dloop1*) in the cytosol of *Srsf6* KD RAW Mφ cells (Fig. 2B). Additionally, we determined the necessity of mtDNA to drive the increased basal ISG expression in *Srsf6* KD macrophages, as mtDNA depletion via treatment with the nucleoside analog 2,3’-dideoxycytidine (ddC) (Fig. 2C), returned expression of the ISG *Rsad2* to that of SCR control cells (Fig. 2D). Elevated basal ISG expression was dependent on cytosolic DNA sensing, as *Srsf6* KD did not cause *Rsad2* transcript accumulation in the absence of the cytosolic DNA sensor, cGAS (Fig. 2E-F, S2A). Together, these data argue that loss of SRSF6 causes leakage of mtDNA into the cytosol where it engages cGAS, leading to phosphorylation of IRF3 (Fig. 1L), expression of IFN-β (Fig 1M), and activation of ISG expression in resting macrophages (Fig. 1A-C).

**Figure 2:**
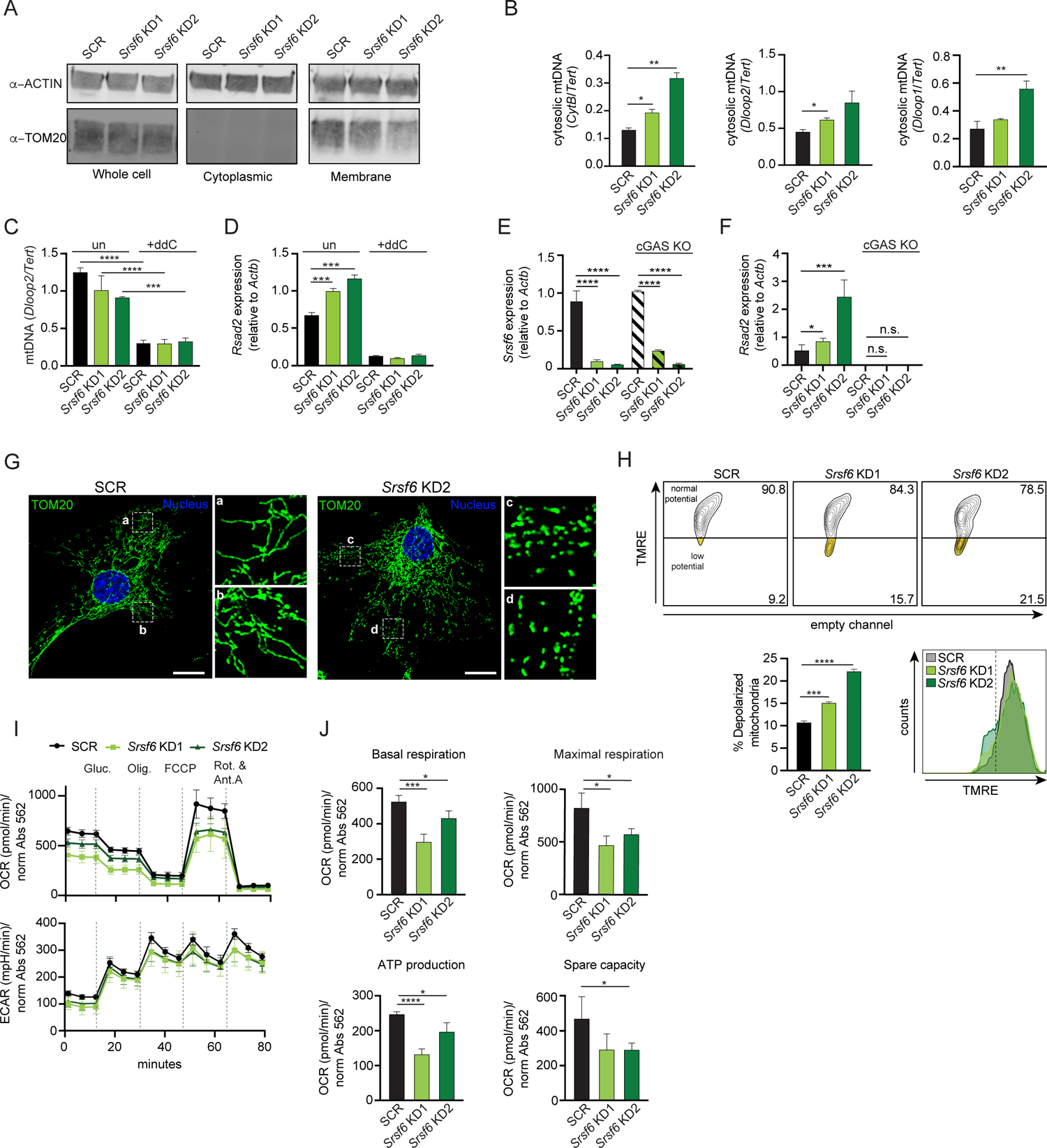
SRSF6 limits cytosolic mtDNA release by maintaining mitochondrial homeostasis. **A.** Immunoblot of mitochondria (TOM20) in total, cytoplasmic, and membrane fractions of *Srsf6* KD RAW 264.7 macrophages. **B.** RT-qPCR of mtDNAs *CytB, Dloop1, Dloop2* relative to nuclear DNA *Tert* in cytosolic fractions of *Srsf6* KD RAW 264.7 cells. **C.** RT-qPCR of total mtDNA *Dloop2* relative to nuclear DNA *Tert* in *Srsf6* KD and SCR control RAW 264.7 cells with or without mtDNA depletion for 8 days. **D.** As in C but measuring *Rsad2*. **E.** RT-qPCR of *Srsf6* in cGAS KO RAW 264.7 cells. **F.** As in E but measuring *Rsad2*. **G.** Immunofluorescence microscopy images visualizing mitochondria in *Srsf6* KD MEFs immunostained with TOM20. Scale bar = 10μM. **H.** Mitochondria membrane potential measured by TMRE staining of *Srsf6* KD RAW 264.7 cells. **I.** Oxygen consumption rate (OCAR) and Extracellular acidification rate (ECAR) measured by Seahorse in *Srsf6* KD RAW 264.7 cells. **J.** Basal respiration, maximal respiration, ATP production, and spare capacity of *Srsf6* KD RAW 264.7 cells determined by OCAR analysis. All data is compared to a scramble control unless indicated. Data are expressed as a mean of three or more biological replicates with error bars depicting SEM. Statistical significance was determined using two tailed unpaired student’s t test. *=p<0.05, **=p<0.01, ***=p<0.001, ****=p<0.0001.

To begin understanding how loss of SRSF6 disrupts mitochondrial homeostasis to allow for mtDNA cytosolic access, we first visualized the mitochondrial network in *Srsf6* KD and control MEFs, which are well-suited for imaging due to their extensive mitochondrial network. Immunofluorescence microscopy (anti-TOM20) revealed increased mitochondrial fragmentation in *Srsf6* KD MEFs (Fig. 2G). Mitochondria in *Srsf6* KD RAW MΦ also exhibited loss of membrane potential (Fig. 2H), as measured by tetramethylrhodamine ethyl ester (TMRE) signal (decreased TMRE signal = increased membrane depolarization). As another measure of mitochondrial function, we measured metabolic output in *Srsf6* KD vs. SCR RAW MΦ cells normalized to protein abundance via the Agilent Seahorse Metabolic Flex Analyzer. *Srsf6* KD cells displayed deficiencies in multiple readouts of oxygen consumption rate (OCR) which serves as a proxy for oxidative phosphorylation (OXPHOS). Loss of SRSF6 reduced mitochondrial OXPHOS indicated by lower basal and maximal respiration. Additionally, *Srsf6* KD mitochondria had lower ATP production, and lower respiratory capacity compared to SCR controls (Fig. 2I top-J). Glycolysis, as measured by extracellular acidification rate (ECAR) was unaffected by SRSF6 knockdown (Fig. 2I, bottom), suggesting that SRSF6 specifically impacts cellular metabolism through mitochondrial OXPHOS.

### SRSF6-dependent alternative splicing of BAX regulates basal ISG expression in macrophages

Our data suggest that SRSF6 limits basal type I IFN expression in macrophages through a role in maintaining mitochondrial homeostasis. SRSF6 is best known as a regulator of alternative splicing (Filippov et al., 2008; Juan-Mateu et al., 2018; Tran and Roesser, 2003; Tranell et al., 2010). Therefore, we hypothesized that loss of SRSF6 could alter alternative splicing of one or more pre-mRNAs that encode proteins involved in mitochondrial biology. To identify SRSF6-dependent alternative splicing events, we mined a list of local splicing variations (LSVs) in *Srsf6* KD MΦ cells quantified by the computational algorithm MAJIQ (Modeling Alternative Junction Inclusion Quantification) (Vaquero-Garcia et al., 2016) as reported in (Wagner et al., 2021). As expected for an SR protein, SRSF6 mostly controls exon inclusion in macrophages (1043 exon skipping LSVs), with changes in intron retention (247 LSVs), and alternative 5’ (206 LSVs) and 3’ (233 LSVs) splice site usage also detected (Fig. 3A). Manual annotation of these >1600 LSVs identified 32 alternatively spliced mitochondrial RNAs, also in the differential expressed gene hits (Fig. 3B). We used semi-quantitative RT-PCR to validate these hits and confirmed that loss of SRSF6 promotes retention of intron 2, in the *Xaf1* pre-mRNA, which is predicted to introduce a premature stop codon and target *Xaf1* for nonsense mediated decay (Fig. S3A).

**Figure 3:**
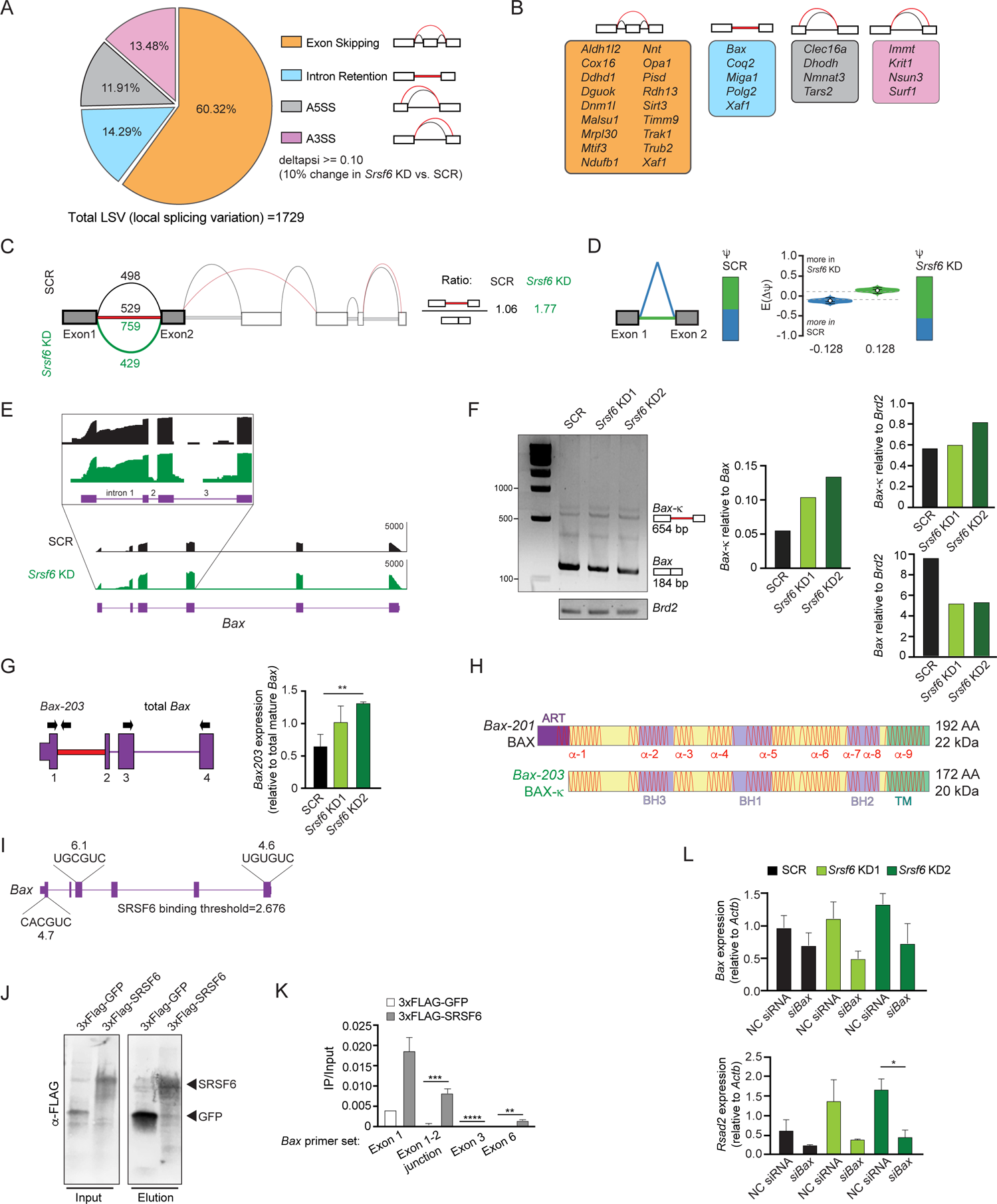
SRSF6 controls alternative splicing of the mitochondrial apoptotic factor BAX. **A.** Percentages of alternative splicing (AS) events in *Srsf6* KD RAW 264.7 macrophages (deltapsi >=0.1). **B.** Categorization of alternative splicing events in mitochondria genes differentially expressed in *Srsf6* KD RAW 264.7 cells. Red lines are AS events and black lines are WT events. **C.** Splice graph of *Bax* in SCR (top) and *Srsf6* KD (bottom) RAW 264.7 cells generated by MAJIQ/VOILA. Intron 1 retention reads relative to exon1-2 junction reads in each genotype shown on right. **D.** MAJIQ Ψ quantification of junctions as illustrated in (C) from SCR (left) and *Srsf6* KD (right) RAW 264.7 cells. Intron retention displayed in green; intron removal displayed in blue. **E.** Integrative Genomics Viewer (IGV) tracks of *Bax,* highlighting exon 1 to exon 3. Zoom-in (top) uses a log scale to facilitate appreciation of the intron reads. **F.** Semi-quantitative RT-PCR of *Bax* and *Brd2* (control) in *Srsf6* KD RAW 264.7 cells with densiometric quantification (LICOR) on right. Gel shown is representative of n>3. **G.** RT-qPCR of *Bax*203 relative to mature *Bax* expression in *Srsf6* KD RAW 264.7 cells. Primers shown on schematic. **H.** Schematics of BAX and BAX-κ proteins. Alpha-helical domains shown as red lines. ART = apoptosis regulatory targeting domain (Goping et al., 2008). **I.** Diagram of predicted *Srsf6* binding sites in *Bax* pre-mRNA with predicted binding strength scores (from ESE Finder). **J.** CLIP Immunoblot of 3xFLAG-GFP and 3xFLAG-SRSF6 constructs expressed in RAW 264.7 cells for 24 h. **K.** CLIP RT-qPCR of 3xFLAG-GFP and 3xFLAG-SRSF6 RT-qPCR of *Bax* exon 1, exon1-2 junction, exon 3, and exon 6. Data shown as IP relative to input. **L.** RT-qPCR of *Bax* and *Rsad2* in *Srsf6* KD RAW 264.7 cells with *Bax* KD via siRNA transfection. All data is compared to a scramble control unless indicated. Data are expressed as a mean of three or more biological replicates with error bars depicting SEM. Statistical significance was determined using two tailed unpaired student’s t test. *=p<0.05, **=p<0.01, ***=p<0.001, ****=p<0.0001.

One alternative splicing event that piqued our interest occurred in *Bax*, the mitochondrial pore-forming protein and executioner of apoptosis. Specifically, MAJIQ analysis measured preferential retention of intron 1 in *Srsf6* KD macrophages (Fig. 3C). Considerable *Bax* intron 1 retention occurred in SCR MΦ cells as well, but to a lesser extent (Fig. 3C-E). We validated SRSF6-dependent *Bax* splicing by semi-quantitative RT-PCR, using a forward primer in exon 1 and a reverse primer in exon 6 (Fig. 3F), as well as by RT-qPCR, using a forward primer in exon 1 and a reverse primer in intron 1 (Fig. 3G). Retention of intron 1 in *Bax* causes usage of a downstream ATG start site encoded in exon 2, which creates a 20 amino acid N-terminal truncation, but otherwise leaves the *B*ax coding sequence completely in-frame and intact (Fig. 3H and S3B). This previously annotated isoform of BAX, dubbed BAX kappa (BAX-κ; GenBank accession number AY095934), is upregulated in the rat hippocampus following cerebral ischemic injury and promotes apoptosis when overexpressed in murine hippocampal neuronal cells (Jin et al., 2001). Because they both lack an ART (<u>a</u>poptosis-regulating targeting) sequence (Cartron et al., 2005; Goping et al., 1998), Bax-κ would be predicted to be functionally analogous to the human BAX-psi isoform, which has been shown to constitutively associate with mitochondria (Cartron et al., 2005). ESE Finder, a web-based platform to identify exonic splicing enhancer motifs (Cartegni et al., 2003), identified three strong SRSF6 consensus sequences in *Bax* exons 1, 3, and 6 (Fig. 3I). UV Crosslinking-immunoprecipitation (CLIP) experiments in RAW MΦ cells showed a clear enrichment of *Bax* transcripts bound to 3xFLAG-SRSF6 compared to a 3xFLAG-GFP control (Fig. 3J-K). Together, these data strongly suggest that SRSF6 controls splicing of *Bax* intron 1 by directly binding consensus exonic splicing enhancers in *Bax* exons.

Previous literature has demonstrated that in addition to promoting release of cytochrome c during apoptosis, BAX pores can release mtDNA capable of stimulating cGAS-dependent type I IFN responses (Rongvaux, 2018; Rongvaux et al., 2014). We hypothesized that high basal ISGs in *Srsf6* KD macrophages are a consequence of Bax-κ-dependent release of mtDNA. Consistent with this prediction, even modest knockdown of *Bax* via siRNA transfection (all isoforms), was capable of rescuing *Rsad2* expression in *Srsf6* KD macrophages (Fig. 3L).

### SRSF6 deficiency sensitizes macrophages to cell death

In addition to their role in cellular metabolism and energy production, mitochondria serve as gatekeepers of multiple cell death pathways. Based on previous studies of BAX-κ and human BAX-psi, we hypothesized that accumulation of BAX-κ would promote cell death in RAW MΦ We first tested this by measuring incorporation of the cell viability stain, propidium iodide (PI) in resting SCR and Srsf6 KD MΦ cells. We observed significantly higher levels of PI incorporation in *Srsf6* KD cell lines (Fig. 4A), suggesting that even in the absence of cell death inducing agents, a population of stable *Srsf6* KD MΦ undergoes programmed cell death. Because *Srsf6* KD MΦ secrete unusually high levels of IFN-β, we tested whether cell death was dependent on IFN-β as in (Apelbaum et al., 2013; Sarhan et al., 2019). Treatment with an anti-IFN-β neutralizing antibody had no effect on PI incorporation in SCR or *Srsf6* KD cells, effectively ruling out a role for IFN−β in mediating SRSF6-dependent cell death (Fig. 4B). Flow cytometric analysis of *Srsf6* KD cells via PI and annexin V staining (a protein that preferentially binds to phosphatidylserine on cells undergoing apoptosis), confirmed higher levels of PI incorporation in the absence of *Srsf6* (Fig. 4C-D)). It also identified a population of pro-apoptotic cells (annexin V-positive but PI-negative) that is more abundant in *Srsf6* KDs (6.3% in KD1 and 9.7% in KD2 vs. 4.2 in SCR) (Fig. 4E). Consistent with *Srsf6* KD RAW 264.7 cells being prone to apoptosis, significantly more *Srsf6* cells stained PI+ after treatment with low levels of the apoptosis-inducing drugs staurosporine (Fig. 4F) or ABT373 (Fig. 4G) and at early time-points following treatment with the cell death agonist etoposide (Fig. S4A).

**Figure 4:**
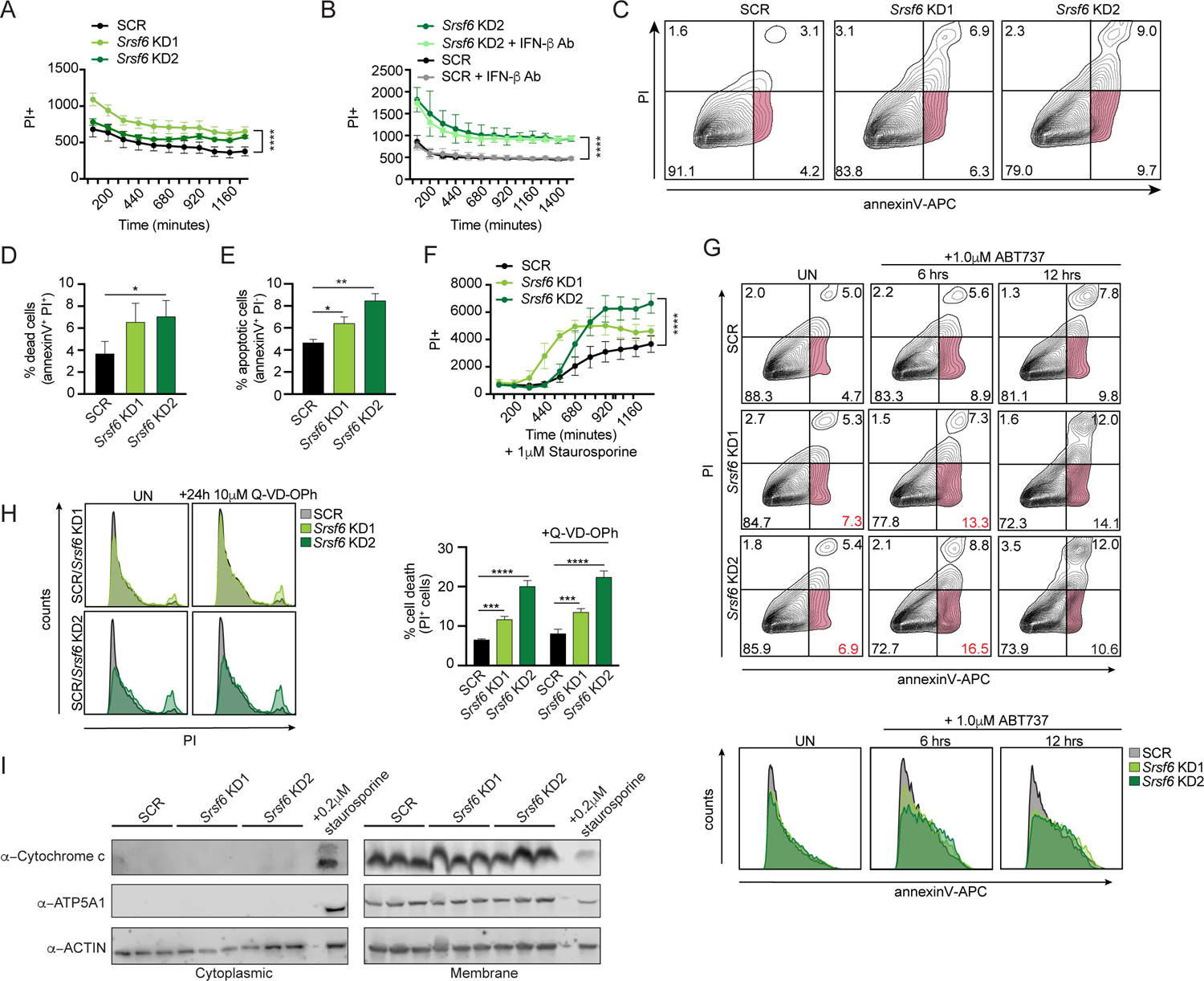
Loss of SRSF6 sensitizes macrophages to caspase-independent apoptotic cell death. **A.** Cell death over a time course in *Srsf6* KD RAW 264.7 cells. **B.** Cell death over a time course in *Srsf6* KD RAW 264.7 cells treated with IFN-β neutralizing antibody. **C.** Apoptotic cell death measured by flow cytometry using APC conjugated annexin V (annexinV-APC) and propidium iodide (PI) dyes in *Srsf6* KD RAW 264.7 cells. **D.** Quantification of dead cells in *Srsf6* KD RAW 264.7 cells from C. **E.** Quantification of apoptotic cells in *Srsf6* KD RAW 264.7 cells from C. **F.** Cell death over a time course in *Srsf6* KD RAW 264.7 cells treated with 1 μM staurosporine. **G.** Apoptotic cell death over a time course measured by flow cytometry using annexinV-APC and PI in *Srsf6* KD RAW 264.7 cells treated with 1 μM ABT737. Histograms display annexinV-APC single stain in *Srsf6* KD. Red numbers indicate annexinV+/PI-cells in *Srsf6* KDs. **H.** Histogram showing cell death after caspase inhibition by flow cytometry in *Srsf6* KD RAW 264.7 cells. Cell death quantification (right). **I.** Immunoblot of cytochrome c in cytoplasmic and membrane fractions of *Srsf6* KD RAW 264.7 cells. SCR cells treated with 0.2 μM staurosporine for 24 h used as a positive control. All data is compared to a scramble control unless indicated. Data are expressed as a mean of three or more biological replicates with error bars depicting SEM. Statistical significance was determined using two tailed unpaired student’s t test. *=p<0.05, **=p<0.01, ***=p<0.001, ****=p<0.0001.

We next set out to better define the nature of cell death in *Srsf6* RAW MΦ cells. Previous reports have implicated BAX in non-canonical proinflammatory forms of cell death and IL-1β release mediated by caspase 8 (Hu et al., 2020; Vince et al., 2018). We did not suspect inflammasome-mediated cell death could explain our findings in *Srsf6* KD RAW MΦ cells as RAW cells lack the inflammasome adapter ASC (Pelegrin et al., 2008). However, given that the cell death observed was not silent, with ISG upregulation, we sought to rule out additional proinflammatory pathways present during *Srsf6*-mediated cell death. Therefore, we knocked down SRSF6 in primary BMDMs and detected no significant difference in IL-1β release between BMDMs transfected with a non-targeting siRNA vs. *Srsf6*-targeting siRNA in either unprimed or poly dA:dT primed cells (Fig. S4B-C). Curiously, despite displaying an apoptotic signature (Fig. 4C, E), cell death in *Srsf6* KD RAW MΦ cells was caspase-independent, as treatment with the pan-caspase inhibitor Q-VD-OPh had no impact on PI incorporation in resting *Srsf6* KD cells (Fig. 4H). Consistent with lack of caspase involvement, resting *Srsf6* KD RAW MΦ did not release cytochrome c (Fig. 4I). These observations, coupled with the disruption to mitochondrial membrane potential reported in (Fig. 2H), suggest a form of mitochondrial-dependent caspase-independent cell death (CICD) (as described by Xiang et al 1996, reviewed in Tait and Green 2008) that occurs preferentially in *Srsf6 KD* macrophages. We hypothesize that BAX-κ, which can trigger mitochondrial outer membrane permeabilization (MOMP) and release of mtDNA without releasing cytochrome c, is a major contributor to this noncanonical form of cell death.

### BAX-κ is sufficient to drive basal ISG expression and apoptosis in macrophages

The results of our *Bax* KD experiment suggest that BAX is necessary to drive type I IFN expression in *Srsf6* KD cells (Fig. 3L). To specifically implicate BAX-κ in this phenotype, we generated C-terminal 2xSTREP-tagged constructs of BAX, BAX-κ, and a BAX point mutant (BAX^G179P^) that is unable to translocate to mitochondria (Kuwana et al., 2020) and introduced them via lentiviral transduction into RAW MΦ cells expressing a doxycycline-inducible transactivator (Fig. S5A). Because high levels of doxycycline can negatively impact mitochondrial function (Dijk et al., 2020), we experimentally determined the lowest doxycycline concentration that ensured robust and uniform expression of an inducible mCherry construct without significant toxicity (1 μg/mL) (Fig. S5A-C). All three of these constructs were expressed in a doxycycline-inducible fashion, albeit to much lower levels than that of a 2xSTREP-GFP control (Fig. 5A). Remarkably, we found that expression of BAX-κ was sufficient to induce robust basal ISG expression in the wild type RAW doxycycline-inducible cell line (Fig. 5B). BAX-κ expression was also sufficient to induce apoptosis and cell death, as measured by flow cytometry at 15 h post-doxycycline treatment (Fig. 5C) or by PI+ incorporation over a 15 h time-course of doxycycline induction (Fig. 5D). Interestingly, unlike the cell death we measured in resting *Srsf6* KD RAW MΦ cells, BAX-κ induced apoptosis and cell death was largely caspase-dependent (Fig. 5C, bottom graphs). We predict that high levels of BAX-κ in these overexpression cell lines induces a level of MOMP that allows for cytochrome c release and downstream caspase-activation. Importantly, stimulation of apoptosis by staurosporine caused significant cell death in both BAX and BAX-κ expressing macrophages, confirming that an activation signal is required for full length BAX to stimulate cell death (Fig. 5E). Collectively, these data strongly argue that BAX-κ accumulation triggers the type I IFN and cell death phenotypes we uncovered in *Srsf6* KD macrophages.

**Figure 5:**
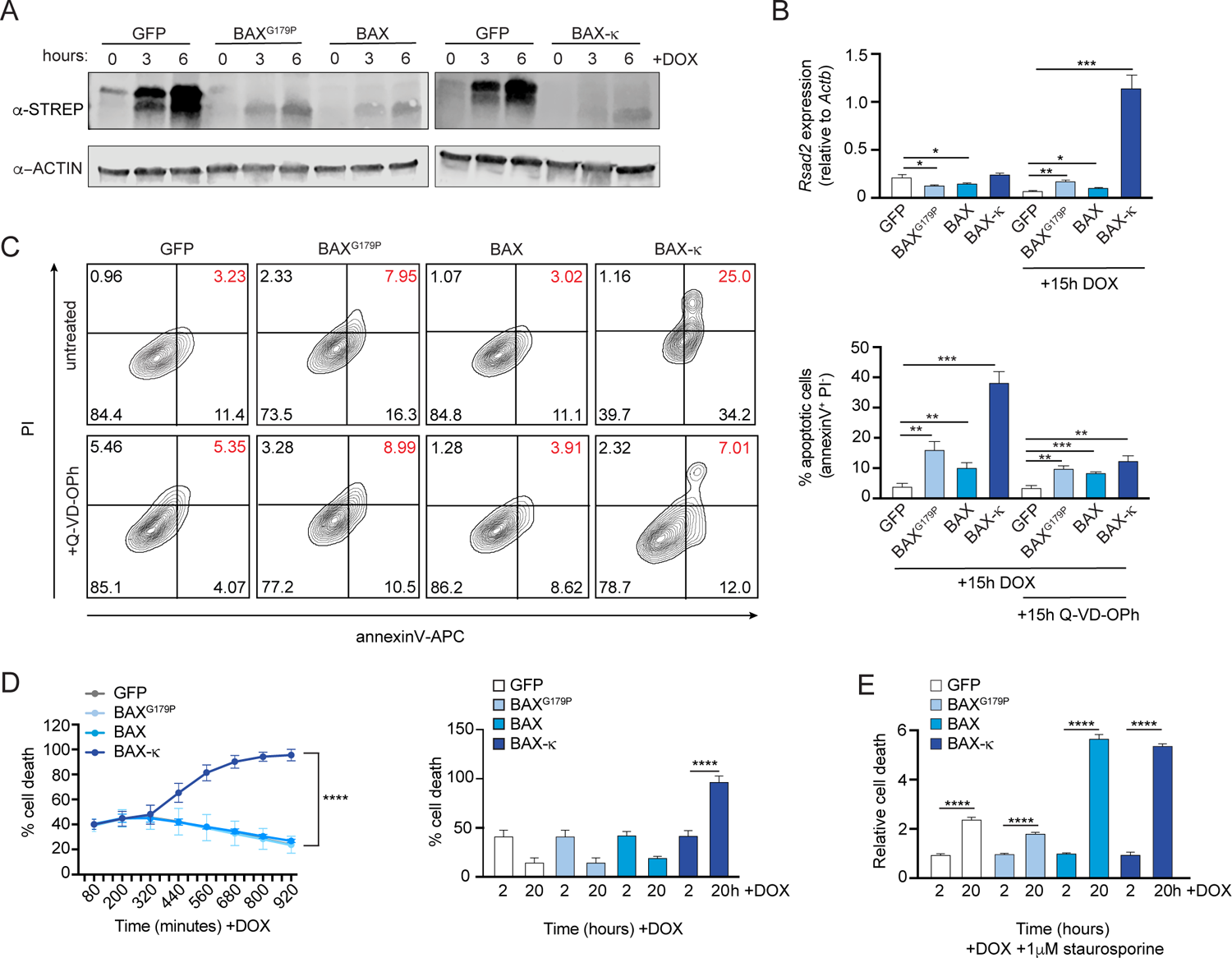
Expression of BAX-κ promotes mtDNA release and cell death in macrophages. **A.** BAX^G179P^, BAX, and BAX-κ inducible RAW 264.7 cells expressed over a time course after addition of doxycycline (DOX). **B.** Expression of *Rsad2* 15 h after DOX induction in GFP, BAX^G179P^, BAX, and BAX-κ-expressing RAW 264.7 cells by RT-qPCR. **C.** Apoptotic cell death measured by flow cytometry using annexinV-APC and PI in GFP, BAX^G179P^, BAX, and BAX-κ inducible macrophages expressed for 15 h with 1 μg DOX and caspase inhibitor (10 μM Q-VD-OPh). Red numbers indicate dead cells (AnnexinV+/PI+). Apoptotic cells (AnnexinV+/PI-) quantification (right). **D.** Cell death over a time course after DOX induced expression of GFP, BAX^G179P^, BAX, and BAX-κ. Starting and ending cell death (PI+) shown as a bar graph on right. **E.** Relative cell death measured by PI incorporation at 2h and 20h after DOX-induced expression of GFP, BAX^G179P^, BAX, and BAX + addition of 1 μM staurosporine. Data are expressed as a mean of three or more biological replicates with error bars depicting SEM. Statistical significance was determined using two tailed unpaired student’s t test. *=p<0.05, **=p<0.01, ***=p<0.001, ****=p<0.0001.

### SRSF6 phosphorylation regulates *Bax* alternative splicing and cell death

Our original interest in SR protein function in macrophages was borne out of a global phosphoproteomics analysis that reported SR proteins, including SRSF6 were differentially phosphorylated in BMDMs at several sites over a 24h time course of infection with the intracellular bacterial pathogen *Mycobacterium tuberculosis* (Mtb) (Budzik et al., 2020). Having discovered a role for SRSF6 in alternative splicing of BAX and regulating programmed cell death, we set out to determine whether phosphorylation of SRSF6 impacted its ability to carry out these activities in macrophages. Leveraging data from Budzik et al., we prioritized three serine residues that were differentially phosphorylated in studies of Mtb-infected (Budzik et al., 2020) or LPS-stimulated macrophages (Weintz et al., 2010): S295, S297, and S303 (Fig. 6A). To test the contribution of phosphorylation of these serines to SRSF6 function, we generated RAW MΦ cell lines expressing doxycycline-inducible constructs of wild-type SRSF6 alongside phosphodead (Ser(S)-to-Ala(A)) or phosphomimetic (Ser(S)-to-Asp(D)) mutations at each site. Expression of each SRSF6 allele was confirmed by immunoblot at 24 h following addition of doxycycline (Fig. 6B). Because phosphorylation of SRSF6 has been implicated in nuclear/cytoplasmic shuttling (Jeong, 2017), we tested where each SRSF6 mutant accumulated in cells using confocal immunofluorescence microscopy, using an antibody directed against the 3xFLAG tag. We observed no major differences in SRSF6 protein localization in any of our phosphomutant/mimetic cell lines compared to wild-type SRSF6 (predominantly nuclear in all cases) (Fig. 6C).

**Figure 6.**
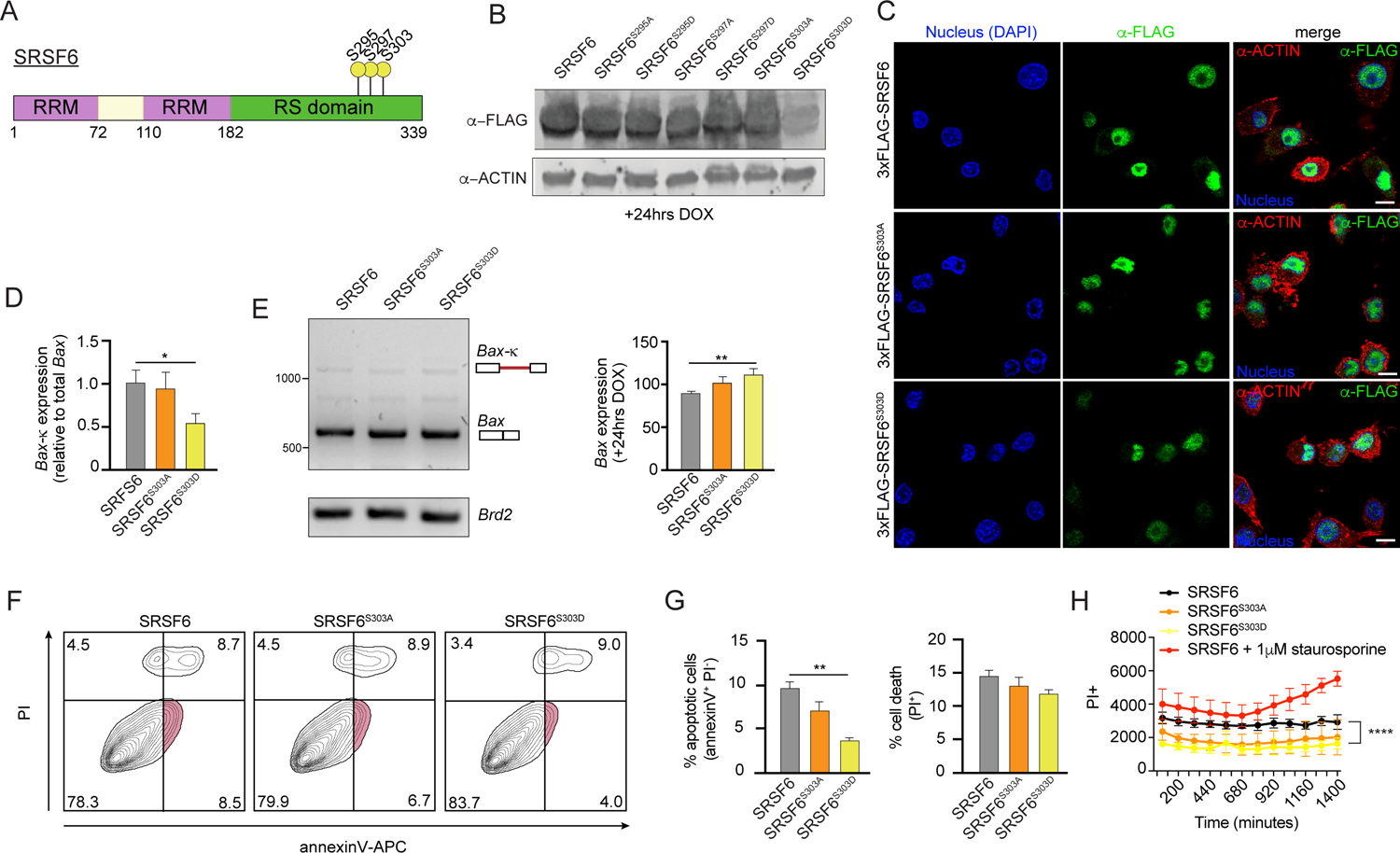
Phosphorylation of SRSF6 at S303 promotes splicing of *Bax* to limit *Bax*-κ expression and prevent cell death. **A.** Diagram of differentially phosphorylated residues in SRSF6 according to Budzik et al., 2020. **B.** Immunoblot of FLAG tagged SRSF6, SRSF6^S295A^, SRSF6^295D^, SRSF6^S297A^, SRSF6^S297D^, SRSF6^S303A^, and SRSF6^S303D^ inducible RAW 264.7 macrophages expressed for 24 h after DOX induction. **C.** Immunofluorescence microscopy images visualizing 3x-FLAG tagged SRSF6, SRSF6^S303A^, and SRSF6^S303D^ inducible RAW 264.7 cells expressed for 24 h after DOX induction. **D.** RT-qPCR of *Bax*203 in FLAG-tagged SRSF6, SRSF6^S303A^, and SRSF6^S303D^ inducible RAW 264.7 cells after DOX induction for 24 h. **E.** RT-PCR of *Bax* and *Brd2* in FLAG-tagged SRSF6, SRSF6^S303A^, and SRSF6^S303D^ inducible RAW 264.7 cells expressed for 24h after DOX induction with quantification of multiple independent experiment. Representative gel shown. **F.** Apoptotic cell death measured by flow cytometry using annexinV-APC and PI in FLAG tagged SRSF6, SRSF6^S303A^, and SRSF6^S303D^ inducible RAW 264.7 Mφ expressed for 24 h after DOX induction. **G.** %apoptotic cells and %dead cells from D. **H** Cell death over a time course in FLAG tagged SRSF6, SRSF6^S303A^, and SRSF6^S303D^ inducible RAW 264.7 cells. FLAG-tagged SRSF6 inducible RAW 264.7 cells were treated with 1 μM staurosporine as a positive control. Data are expressed as a mean of three or more biological replicates with error bars depicting SEM. Statistical significance was determined using two tailed unpaired student’s t test. *=p<0.05, **=p<0.01, ***=p<0.001, ****=p<0.0001.

To begin to implicate specific phosphorylation events in SRSF6 activity, we measured the ratio of *Bax-*κ relative to total *Bax*, as in Fig. 3G, in each of the SRSF6-expressing cell lines. We observed that expression of SRSF6^S303D^ decreased levels of *Bax-*κ transcripts, compared to those in wild-type SRSF6-expressing macrophages (Fig. S6A and 6D). Likewise, SRSF6^S303D^-expressing cell lines had higher levels of canonical *Bax* transcripts, relative to wild-type SRSF6-expressing cells, by semi-quantitative RT-PCR (Fig 6E). Consistent with altered Bax-κ to Bax ratios in SRSF6^S303D^-expressing cell lines, expression of the SRSF6^S303D^ phosphomimetic allele rendered cells less prone to apoptosis (annexin V+/PI-) and cell death (Fig. 6F-H). Together, these data suggest that phosphorylation of SRSF6 at S303 promotes splicing of *Bax*, to generate the canonical BAX protein and limit apoptosis.

### Regulation of SRSF6 controls infection outcomes in macrophages

Our data support a model wherein SRSF6 maintains cellular homeostasis and basal type I IFN expression by controlling the abundance of BAX isoforms. Therefore, we predicted that expression of SRSF6 itself could be subject to regulation downstream of pathogen sensing, as a way for cells to prime apoptotic cell death and/or cytosolic DNA sensing via mtDNA release. To further appreciate how macrophages regulate SRSF6 activity, we measured *Srsf6* transcript levels during infection and in response to immune agonists. We consistently detected an approximately 2-fold decrease in *Srsf6* transcript abundance at various time points following *M. tuberculosis* infection of macrophages, as well as in the lung homogenates collected from Mtb-infected mice (day 21 and 77 post-infection) (Fig. 7A-B). Macrophages sense Mtb infection via several pattern recognition receptors, including dsDNA via cGAS and autocrine sensing of IFN-β via IFNAR (Watson et al., 2015). Direct stimulation of these pathways via transfection of dsDNA (Fig. 7C) or treatment with recombinant IFN-β (rIFN-β) (Fig. 7D) also led to downregulation of *Srsf6* transcript abundance, as did infection with the gram-negative bacterial pathogen *Salmonella enterica* serovar Typhimurium (Fig. 7E), engagement of TLR-4 via LPS (Fig. 7F) and infection with the RNA virus VSV (Fig. 7G). These data suggest that downregulation of *Srsf6* is part of the general macrophage pathogen sensing response.

**Figure 7:**
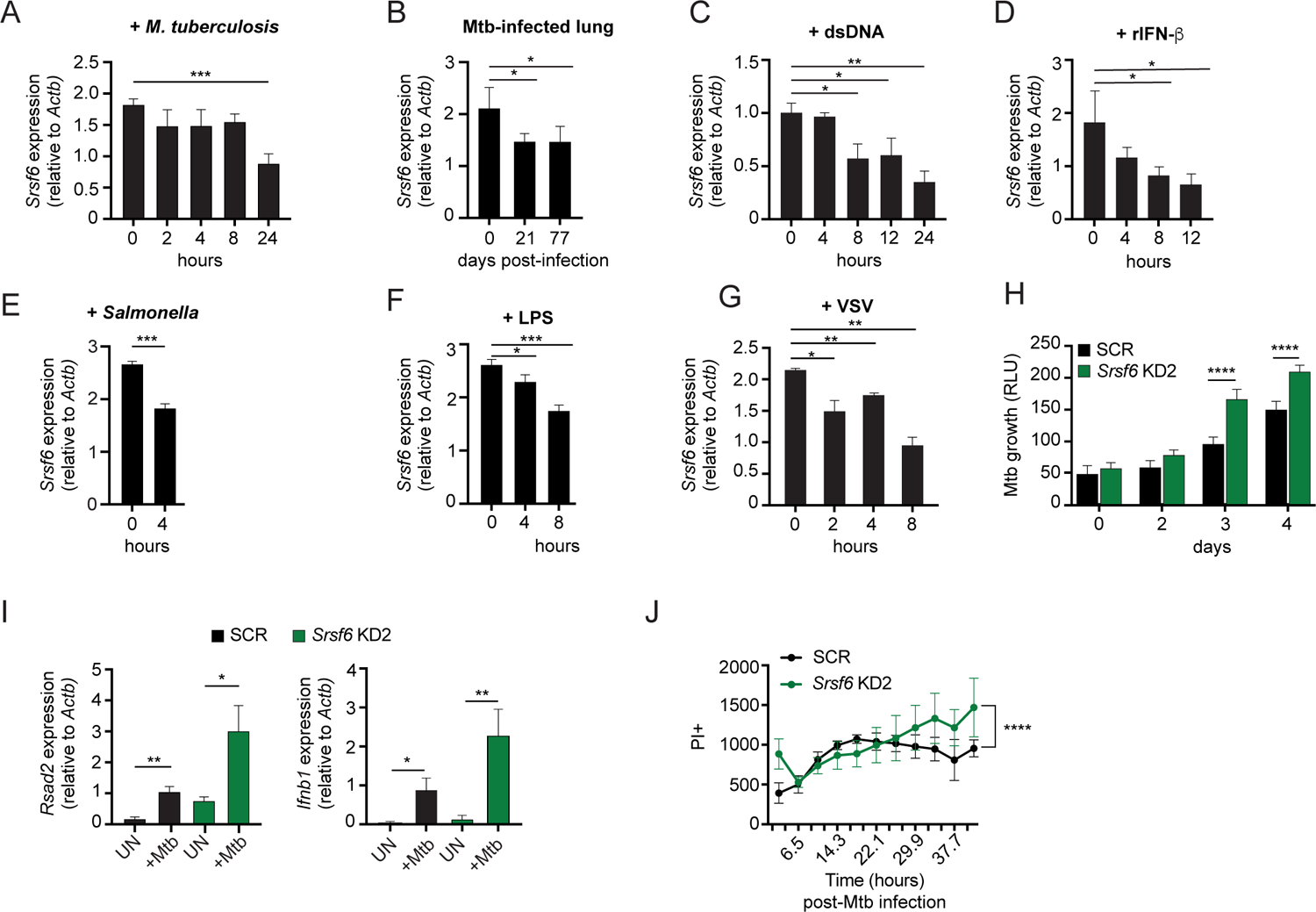
Modulation of SRSF6 expression contributes to innate immune control of the intracellular bacterial pathogen *Mycobacterium tuberculosis*. **A.** RT-qPCR of *Srsf6* in RAW 264.7 macrophages infected with *Mycobacterium tuberculosis* (Mtb) (MOI=5) over a time course. **B.** RT-qPCR of *Srsf6* in Mtb-infected mouse lung samples over a time course of *in vivo* infection. **C.** RT-qPCR of *Srsf6* in RAW 264.7 cells treated with 1μg double stranded DNA (dsDNA). over a time course. **D.** As in C but treated with recombinant IFN-β (rIFN-β). **E.** RT-qPCR of *Srsf6* in S. Typhimurium infected RAW 264.7 MΦ (MOI=5) at 0 and 4h. **F.** As in C but treated with LPS. **G.** RT-qPCR of *Srsf6* in VSV infected RAW 264.7 cells (MOI=1) over a time course. **H.** Mtb *luxBCADE* growth in *Srsf6* KD RAW 264.7 cells measured by relative light units (RLUs) over a time course (MOI=1). **I.** RT-qPCR of *Rsad2* and *Ifnb1* in *Srsf6* KD RAW 264.7 cells infected with Mtb at (MOI=10), 4h post-infection. **J.** Cell death over a time course in SCR and *Srsf6* KD RAW 264.7 cells infected with Mtb at (MOI=5). All data is compared to a scramble control unless indicated. Data are expressed as a mean of three or more biological replicates with error bars depicting SEM. Statistical significance was determined using two tailed unpaired student’s t test. *=p<0.05, **=p<0.01, ***=p<0.001, ****=p<0.0001.

Lastly, we set out to investigate whether SRSF6 plays a more direct role in dictating the outcome of infection with an intracellular pathogen like Mtb. To this end, we infected monolayers of SCR and *Srsf6* KD RAW MΦ cells with a luciferase-expressing strain of Erdman Mtb (MOI = 1) as in (Bell et al., 2021; Hoffpauir et al., 2020) and monitored relative light units (RLU) as a measurement of Mtb replication. Remarkably, we found that Mtb grew significantly better in *Srsf6* KD cell lines compared to SCR controls (Fig. 7H). This increase in Mtb replication was concomitant with hyperinduction of *Ifnb1* and interferon stimulated genes, consistent with high basal type I IFN potentiating type I IFN responses (Fig. 7I) (West et al., 2015). Along with harboring higher Mtb bacterial burdens, more *Srsf6* KD RAW MΦ cells stained PI+ compared to SCR controls over a 40+h Mtb infection (Fig. 7J). Collectively, these results demonstrate a critical role for SRSF6 in controlling inflammatory gene expression and cell death during bacterial infection and suggest that regulation of SRSF6, both at the post-translational and transcriptional levels, constitutes an unappreciated layer of complexity in the macrophage innate immune response.

## DISCUSSION

While advances in bioinformatics have facilitated computational detection of alternatively spliced transcripts, assigning function to individual protein isoforms largely remains uncharted territory. Here, we identify the splicing factor SRSF6 as a primary regulator of *Bax* pre-mRNA splicing in murine macrophages. We show that SRSF6-dependent control of the ratio of BAX to BAX-κ isoforms balances critical aspects of innate immune homeostasis in both macrophage-like cell lines and primary macrophages. Specifically, we demonstrate that expression of BAX-κ promotes upregulation of basal type I IFN expression driven by cytosolic mtDNA and renders cells prone to apoptotic cell death. Unlike BAX, which requires a signal to direct mitochondrial targeting and pore formation, BAX-κ is sufficient to trigger type I IFN expression and cell death in resting macrophages (Fig. 5B-D). These findings position BAX-κ as a potential uncoupler of immunostimulatory mitochondrial DAMP release from cell death and suggests that BAX-κ plays a role in promoting the primed antiviral state that innate immune cells, such as macrophages, must maintain.

Connections between SRSF6 and alternative splicing of the cell death protein BAX complement multiple studies that correlate altered SRSF6 expression levels with cancer progression. Our data show that in macrophages, loss of SRSF6 upregulates the pro-apoptotic BAX-κ isoform suggesting that under normal physiological conditions, SRSF6 is a negative regulator of cell death. Consistent with this, upregulation of SRSF6 promotes proliferation of breast cancer cells (Park et al., 2019), colorectal cancer cells (Wan et al., 2019), and lung cancer cells (Cohen-Eliav et al., 2013). Our finding that loss of SRSF6 expression impacts homeostatic levels of IFN-β and ISGs motivates future studies of SRSF6-dependent immune dysregulation in cancer, particularly in cancers of myeloid cells. Perhaps dysregulation of the immune milieu is part of why SRSF6 upregulation is associated with poor cancer outcomes. At the same time, SRSF6 can also promote expression of pro-apoptotic forms of proteins like BIM (BIM-S) (Hara et al., 2013) and cassette exon inclusion in the apoptotic factor FAS (Choi et al., 2022). These seemingly contradictory roles for SRSF6 in both limiting and promoting apoptotic cell death suggest that its activity could be concentration and/or cell-type dependent.

Our report that phosphorylation of SRSF6 at S303 promotes *Bax* splicing and limits BAX-κ-dependent apoptotic cell death adds to a growing literature of regulation of gene expression via post-translational modification of splicing factors. Zn(2+)-dependent phosphorylation of SRSF6 has previously been associated with apoptosis via generation of BIM-S (Hara et al., 2013) and ubiquitin-mediated control of SRSF6 protein levels has been linked to exon skipping in T cell acute lymphoblastic leukemia (Zhou et al., 2020). Our data support a model whereby SRSF6 is regulated at multiple levels downstream of pathogen sensing, through decreasing transcript abundance (Fig. 7) and differential phosphorylation (Budzik et al., 2020). These levels of regulation argue that SRSF6 is a bone fide player in the macrophage innate immune response acting as a common signaling molecule to promote a state of antiviral readiness while regulating apoptotic vs. pro-inflammatory cell death. Determining the precise signals that trigger the regulation of SRSF6 in the nucleus downstream of pattern recognition receptors in the plasma membrane and cytosol remain important future questions.

Seminal work from White et al. and Rongveux et al. implicates apoptotic caspases in suppressing cGAS-dependent type I IFN expression via mtDNA released by BAX/BAK pores during mitochondrial apoptosis (Rongvaux et al., 2014; White et al., 2014). One interpretation of their findings is that cells prevent aberrant type I IFN expression by dying; thus, one cannot capture *Ifnb1* expression triggered by BAX/BAK pore formation without inhibiting or genetically ablating caspases. However, our work and that of others, begin to suggest that BAX/BAK release of mtDNA and apoptotic cell death can in fact be uncoupled (Riley et al., 2018). While our RAW 264.7 macrophage cell lines are selected for stable KD of SRSF6, only a low percentage stain PI+ as cells are grown in tissue culture (approximately 10-20%) (Fig. 4). This suggests that some threshold of BAX-κ expression needs to be reached before cells undergo apoptotic cell death; this threshold is likely maintained by SRSF6, as our reported phenotypes are tightly correlated with the degree of *Srsf6* KD (Fig. 1F-G). Our data also show that cell death in *Srsf6* KD macrophages is caspase-independent and is not concomitant with cytochrome c release (Fig. 4H-I). While additional experiments at the single cell level are needed to confirm that *Srsf6* KD cells can continuously release mtDNA without releasing cytochrome c and triggering apoptosis, the fact that we were able to uncover the *Srsf6* KD basal type I IFN phenotype in the absence of caspase inhibition argues that some degree of uncoupling is indeed at play.

Consequently, regulation of BAX/BAK pore formation may constitute an unappreciated way for cells to maintain homeostatic levels of IFN signaling, prime antiviral responses, and regulate cell death programs. Indeed, there is growing appreciation that the nature of BAX/BAK pores dictates their ability to release mtDNA and trigger cytosolic DNA sensing. Large BAX/BAK “macropores” that form at late stages of apoptosis have been shown to allow herniation and extrusion of the inner membrane, releasing mtDNA from the matrix (McArthur et al., 2018). The availability of BAX and BAK and the composition of BAX/BAK pores (BAX vs. BAK vs. BAX/BAK) also manages the rate at which pores extrude mtDNA. Specifically, BAK pores are more immunogenic than BAX pores and enable faster mtDNA release (Cosentino et al., 2022). It is tempting to speculate BAX-κ promotes mtDNA release and type I IFN expression by disrupting the balance of BAX and BAK at the mitochondria. This could occur by BAX-κ binding to and sequestering BAX away from BAK, allowing for more BAK pore formation. Alternatively, BAX-κ itself could form pores with BAX and/or BAK that preferentially release mtDNA or BAX-κ could form homo-oliogomeric pores with their own unique properties. Detailed visualization of BAX/BAK pores and mtDNA extrusion in cells genetically engineered to express a single BAX isoform (BAX or BAX-κ) alongside biochemical studies of BAX-κ oligomerization will provide important insights into the properties of this isoform and how it contributes to the innate immune phenotypes we report here.

Our finding that the intracellular bacterial pathogen *Mycobacterium tuberculosis* replicates more efficiently in *Srsf6* KD macrophages suggests that maintaining the balance of BAX vs. BAX-κ is needed to restrain bacterial replication and spread. Numerous reports demonstrate that cell death modality usage is a critical factor in determining Mtb pathogenesis. The established paradigm in the Mtb field asserts that apoptosis controls Mtb replication and spread while necrotic cell death promotes it (Abarca-Rojano et al., 2003; Abebe et al., 2011; Aguilo et al., 2013; Aguilo et al., 2014; Behar et al., 2010; Behar et al., 2011). It is curious then, that we observe more Mtb replication and more cell death in *Srsf6* KD RAW MΦ (Fig. 7H and J). It is possible that SRSF6 contributes to Mtb restriction in BAX-independent ways. It is also possible that basal IFN expression in *Srsf6* KD RAW MΦ creates an intracellular milieu that is ill-adapted for destroying intracellular bacteria (but highly efficient at restricting viral replication (Fig. 1P)). Further studies into the precise nature of the cell death that occurs in Mtb-infected *Srsf6* KD RAW MΦ and its reliance on *Ifnb1* may provide important insights into how BAX isoform usage impacts cell-intrinsic control and cell-to-cell spread of pathogens like Mtb. Collectively, this work highlights the role of pre-mRNA splicing in shaping the macrophage proteome and demonstrates how disruption of protein isoform stoichiometry can impact mitochondrial homeostasis and the ability of cells to respond to infection.

## ACKNOWLEDGEMENTS

We would like to thank Dr. A. Phillip West and members of the West lab for providing us with VSV stocks and sharing viral infection protocols. We thank Drs. Jeffery Cox, Bennett Penn, and Jonathan Budzik for generously sharing their Mtb phosphoproteomics datasets with us. We thank Dr. Malea Murphy (TAMU COM Integrated Microscopy and Imaging Laboratory) and Robbie Moore (COM-CAF) and the core facilities for their technical assistance and advice. Finally, we would like to thank the members of the Patrick and Watson laboratories for their support and reviews of the manuscript. This work was funded through NIH/NIGMS R35GM133720 (to KLP) NIH/NIAID R01AI155621 (to ROW).

## SUPPLEMENTAL FIGURES

**Figure S1.**
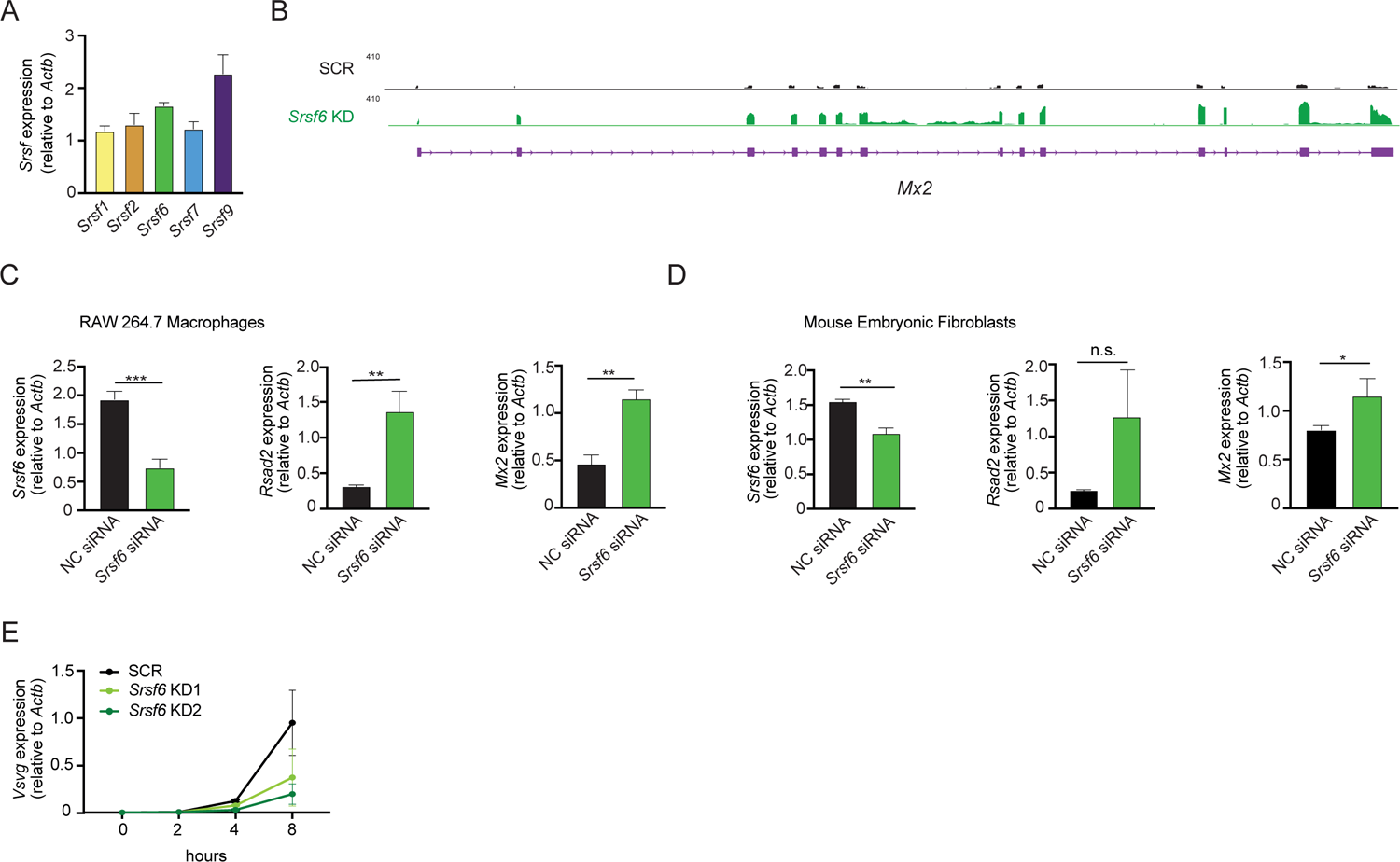
Related to Figure 1. **A.** RT-qPCR of *Srsf* in RAW 264.7 macrophages. **B.** Integrative Genomics Viewer (IGV) tracks of *Mx2* in *Srsf6* KD RAW 264.7 cells. **C.** RT-qPCR of *Srsf6, Rsad2,* and *Mx2* in *Srsf6* siRNA KD RAW 264.7 cells**. D.** As in C but in MEFs. **E.** VSV replication in *Srsf6* KD RAW 264.7 cells at 0, 2, 4, 8 h post infection (MOI=1) measured by RT-qPCR of *Vsvg.* All data is compared to a scramble control unless indicated. Data are expressed as a mean of three or more biological replicates with error bars depicting SEM. Statistical significance was determined using two tailed unpaired student’s t test. *=p<0.05, **=p<0.01, ***=p<0.001, ****=p<0.0001.

**Figure S2.**
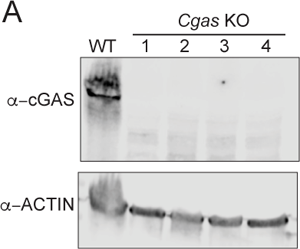
Related to Figure 2. **A.** Immunoblot of cGAS in WT and *cGAS* KO RAW 264.7 cells. cGAS lanes are from multiple protein preparations. Data are expressed as a mean of three or more biological replicates with error bars depicting SEM. Statistical significance was determined using two tailed unpaired student’s t test. *=p<0.05, **=p<0.01, ***=p<0.001, ****=p<0.0001.

**Figure S3.**
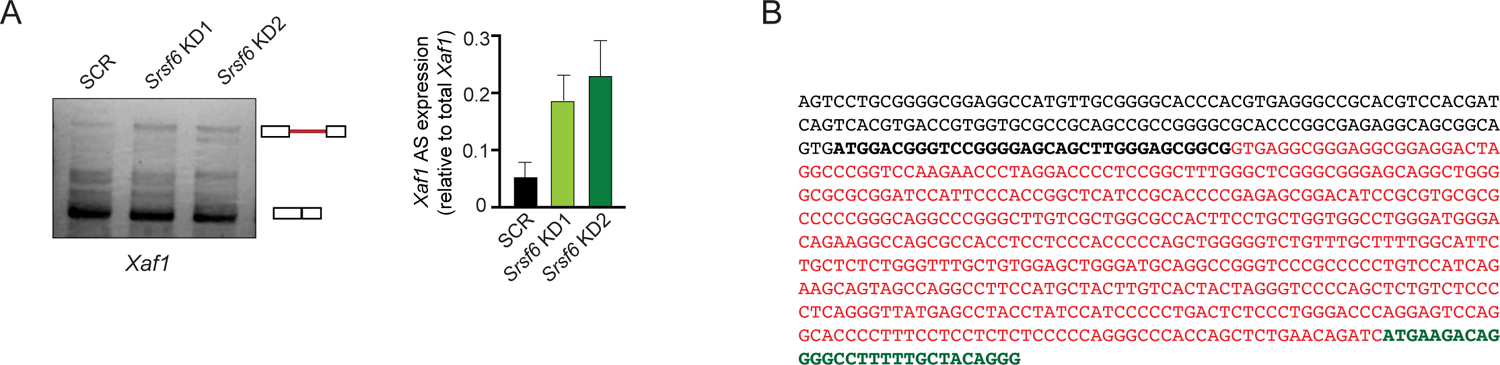
Related to Figure 3. **A.** PCR of *Xaf1* and *Brd2* in *Srsf6* KD RAW 264.7 cells with quantification. **B.** mRNA sequence of *Bax*201 with *Bax*203 truncated isoform (red). All data is compared to a scramble control unless indicated. Data are expressed as a mean of three or more biological replicates with error bars depicting SEM. Statistical significance was determined using two tailed unpaired student’s t test. *=p<0.05, **=p<0.01, ***=p<0.001, ****=p<0.0001.

**Figure S4.**
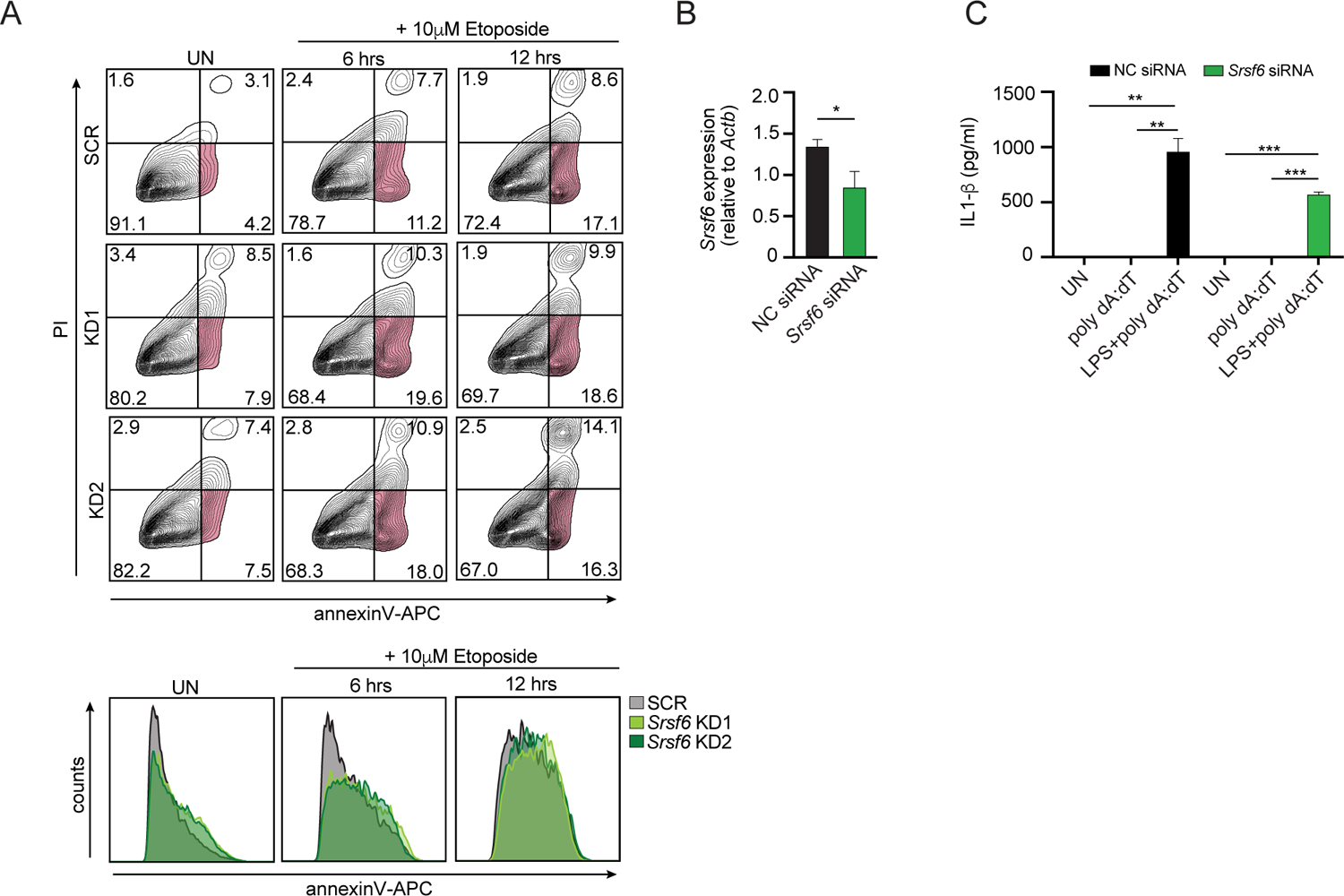
Related to Figure 4. **A.** Apoptotic cell death over a time course measured by flow cytometry using annexinV-APC and PI in *Srsf6* KD RAW 264.7 cells treated with 10 μM etoposide. Histograms display annexinV-APC single stain in *Srsf6* KD RAW 264.7 cells. **B.** RT-qPCR of *Srsf6* in *Srsf6* siRNA KD BMDMs. **C.** Extracellular IL-1β in negative siRNA control and *Srsf6* KD BMDMs untreated and inflammasome treated with LPS 3 h, poly dA:dT 4 h by ELISA with AIM2 inflammasome stimulated positive control (LPS/ poly dA:dT). All data is compared to a scramble control unless indicated. Data are expressed as a mean of three or more biological replicates with error bars depicting SEM. Statistical significance was determined using two tailed unpaired student’s t test. *=p<0.05, **=p<0.01, ***=p<0.001, ****=p<0.0001.

**Figure S5.**
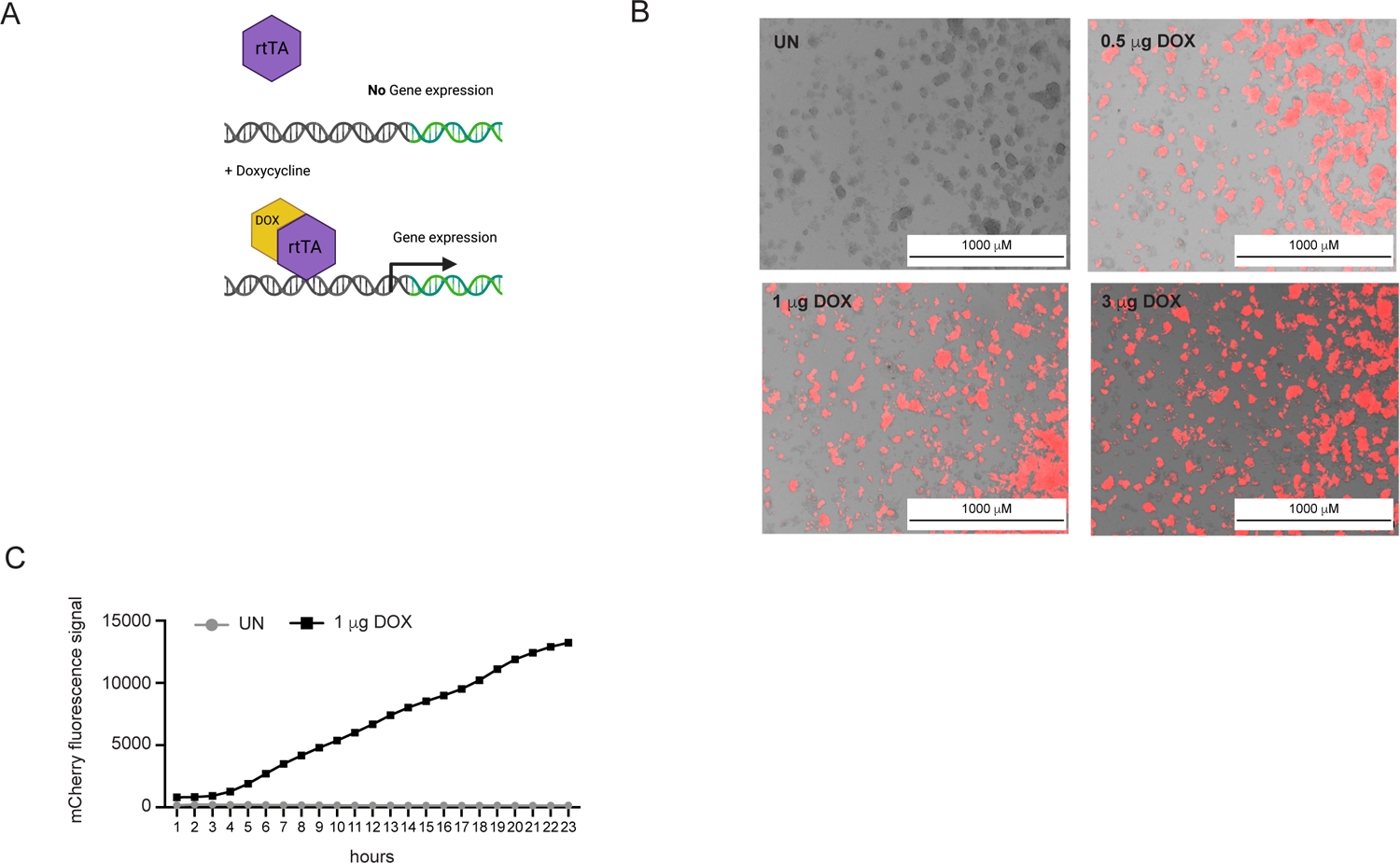
Related to Figure 5. **A.** Schematic of DOX activation of transactivator to induce construct expression **B.** Immunofluorescence/DIC microscopy images visualizing mCherry doxycycline-inducible RAW 264.7 macrophages stimulated with 0.5 μg, 1.0 μg, and 3.0 μg of DOX for 15 h. **C.** mCherry fluorescence over a time course measured using a Lionheart XF analyzer +/- 1.0 μg DOX. Data are expressed as a mean of three or more biological replicates with error bars depicting SEM. Statistical significance was determined using two tailed unpaired student’s t test. *=p<0.05, **=p<0.01, ***=p<0.001, ****=p<0.0001.

**Figure S6.**
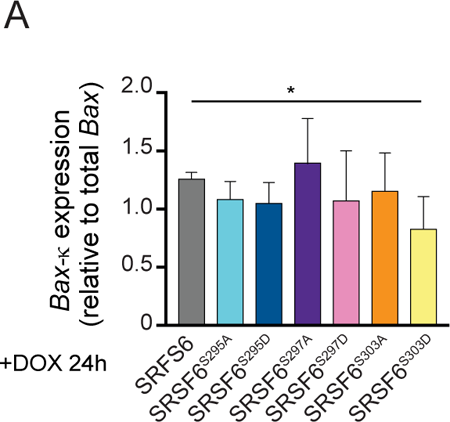
Related to Figure 6. **A.** RT-qPCR of *Bax*203 in FLAG tagged SRSF6, SRSF6^S295A^, SRSF6^S295D^, SRSF6^S297A^, SRSF6^S297D^, SRSF6^S303A^, and SRSF6^S303D^ doxycycline-inducible RAW 264.7 macrophages expressed for 24 h after DOX induction. Data are expressed as a mean of three or more biological replicates with error bars depicting SEM. Statistical significance was determined using two tailed unpaired student’s t test. *=p<0.05, **=p<0.01, ***=p<0.001, ****=p<0.0001.

### EXPERIMENTAL MODELS AND SUBJECT DETAILS

#### M. tuberculosis

The Erdman strain was used for all *M. tuberculosis* (Mtb) infections as well as a luciferase expressing strain. Low passage lab stocks were thawed for each experiment to ensure virulence was preserved. Mtb was cultured in roller bottles at 37 °C in Middlebrook 7H9 broth (BD Biosciences) supplemented with 10% OADC (BD Biosciences), 0.5% glycerol (Fisher), and 0.1% Tween-80 (Fisher). All work with Mtb was performed under Biosafety level 3 containment using procedures approved by the Texas A&M University Institutional Biosafety Committee.

#### *S. enterica* (ser. Typhimirium)

*Salmonella enterica* serovar Typhimurium (SL1344) was obtained from Dr. Denise Monack at Stanford University. S. T. stocks were streaked out on LB agar plates and incubated at 37 °C overnight.

#### Vesicular stomatitis virus

Recombinant Vesicular stomatitis virus (VSV; Indiana serotype) containing a GFP reporter cloned downstream of the VSV G-glycoprotein (VSV-G/GFP) was originally obtained from Dr. John Rose at Yale School of Medicine and shared with us by Dr. A. Phillip West at Texas A&M Health Science Center.

#### M. musculus

Mice used in this study were C57BL6/J (Stock #00064) initially purchased from Jackson labs and afterward maintained with filial breeding. All mice used in experiments were compared to age- and sex-matched controls. Littermates were used for experiments. Mice used to generate BMDMs were males between 10-16 weeks old. For in vivo infection male and female mice were infected with Mtb at 10-12 weeks. Embryos used to make primary MEFs were 14.5 days post-coitum. All animals were housed, bred, and studied at Texas A&M Health Science Center under approved Institutional Care and Use Committee guidelines. All experiments for this study were reviewed and approved by the Texas A&M University Institutional Animal Care and Use Committee (AUP# 2019–0083). Mice were fed 4% standard chow and were kept on a 12 h light/dark cycle and provided food and water ad libitum. Mice were group housed (maximum 5 per cage) by sex on ventilated racks in temperature-controlled rooms.

#### Primary cell culture

Bone marrow derived macrophages (BMDMs) were differentiated from bone marrow (BM) cells isolated by washing mouse femurs with 10 mL DMEM 1 mM sodium pyruvate. Cells were then centrifuged for 5 min at 400 rcf and resuspended in BMDM media (DMEM, 20% FBS (Millipore), 1 mM sodium pyruvate (Lonza), 10% MCSF conditioned media (Watson lab). BM cells were counted and plated at 5×106 cells per 15 cm non-tissue culture treated dishes in 30 mL complete BMDM media. Cells were fed with an additional 15 mL of BMDM media on day 3. Cells were harvested on day 7 with 1X PBS EDTA (Lonza).

Mouse embryonic fibroblasts (MEFs) were isolated from embryos. Briefly, embryos were dissected from yolk sacs, washed 2 times with cold 1X PBS, decapitated, and peritoneal contents were removed. Headless embryos were disaggregated in cold 0.05% trypsin-EDTA (Lonza) and incubated on ice for 20 min., followed by incubation at 37 °C for an additional 20 min. Cells were then DNase treated with 4 mL disaggregation media (DMEM, 10% FBS, 100 μg/mL DNASE I (Worthington)) for 20 min at 37 °C. Isolated supernatants were spun down at 1000 rpm for 5 min. Cells were resuspended in complete MEF media (DMEM, 10% FBS, 1 mM sodium pyruvate), and plated in 15 cm tissue culture-treated dishes 1 dish per embryo in 25 mL of media. MEFs were allowed to expand for 2-3 days before harvest with 0.05% trypsin-EDTA (Lonza).

#### Cell culture

RAW 264.7 macrophages (TIB-71™ ATCC) (originally isolated from male BALB/c mice), L929 ISRE, cGAS knockout RAW 264.7 macrophages, and Lenti-X cells were cultured at 37 °C with a humidified atmosphere of 5% CO2 in complete media containing high glucose, DMEM (Thermo Fisher) with 10% FBS (Millipore) 0.2% HEPES (Thermo Fisher).

#### shRNA knockdowns

For RAW 264.7 macrophages stably expressing scramble knockdown and Srsf6 knockdown, Lenti-X cells were transfected with a pSICO scramble non-targeting shRNA construct and pSICO *Srsf6* shRNA constructs targeted at exon 3 and exon 4 of *Srsf6* using polyjet (SignaGen Laboratories). Virus was collected 24 and 48 h post transfection. RAW 264.7 macrophages were transduced using lipofectamine 2000 (Thermo Fischer). After 48 h, media was supplemented with hygromycin (Invitrogen) to select for cells containing the shRNA plasmid.

#### siRNA knockdowns

Knockdown of mRNA transcripts was performed by plating 3×105 RAW 264.7 macrophages, 3.5×105 BMDMs on day 4 of differentiation, or 2×105 MEFs, in 12-well plates and rested overnight. The following day complete media was replaced with 500 μL fresh complete media 30 min prior to transfection. Cells were transfected using Fugene SI reagent or Viromer Blue reagent and 10 μM of siRNA stock against either Srsf6 (4390771, IDS86053) or Bax (AM16708, ID100458). For a negative control, Silencer® Select Negative Control #1 (4390843) was used. Cells were incubated for 48-72 h in transfection media at 37 °C with 5% CO2 prior to downstream experiments.

#### Doxycycline inducible cell line generation

For generation of doxycycline inducible RAW 264.7 macrophages, pLenti CMV rtTA3 Blast (Addgene w756-1) stably expressing clonal RAW 264.7 macrophages were transduced with pLenti CMV Puro DEST (Addgene w118-1) constructs containing GFP-Strep, Bax201-Strep, Bax203-Strep, BaxG179P-Strep, GFP-FL, SRSF6-FL, and all SRSF6-FL phosphorylation mutants. After 48 h construct containing cells were selected through addition of puromycin (Invivogen). 1 mg/mL doxycycline (Sigma-Aldrich) treatment was used to activate construct expression.

#### *In vitro* infections

##### Mtb infections

To prepare the inoculum, bacteria were grown to mid log phase (OD 0.6–0.8), spun at low speed (500 rcf) to remove clumps, and then pelleted and washed with PBS 2X. Resuspended bacteria were sonicated and spun at low speed again to further remove clumps. The Mtb was then diluted in DMEM plus 10% horse serum (Gibco) and added to cells at a multiplicity of infection (MOI) of 10 for RNA and protein analysis, a MOI of 5 for cell death studies, and a MOI of 1 for bacterial growth assays. The day before the infection, BMDMs were seeded at 3×105 cells per well (12-well dish), and RAW 264.7 macrophages were plated on 12-well tissue culture–treated plates at a density of 3×105 cells per well or plated in corning 96 well black plates at 2.5×104 cells/well and allowed to rest overnight. Cells were spun with bacteria for 10 min at 1,000 rcf to synchronize infection, washed 2X with PBS, and then incubated in fresh media. Where applicable, RNA was harvested from infected cells using 0.5 mL TRIzol reagent at each time point. Protein lysates were harvested with 1X RIPA lysis buffer and boiled for 10 min. For cell death studies propidium iodide (PI) (Invitrogen) was added to media and PI incorporation was measured by fluorescence over time using Lionheart XF analyzer. For bacterial growth assays RAW 264.7 macrophages were plated on 12-well tissue culture–treated plates at a density of 2.5×105 cells per well. Luminescence was read for *M. tuberculosis* luxBCADE by lysing in 250 μl 0.5% Triton X-100 and dividing sample into duplicate wells of a 96-well white bottomed plate (Costar). Luminescence was measured and normalized to background using the luminescence feature of the INFINITE 200 PRO (Tecan) at 0, 48, 72 and 92 h post infection.

#### *S. enterica* (ser. Typhimirium) infections

For S.e Typhimurium infection overnight cultures of bacteria were grown in LB broth containing 0.3 M NaCl and grown at 37 °C until they reached an OD600 of approximately 0.9 RAW 264.7 macrophages were seeded in 12-well tissue culture-treated plates at a density of 7×105 cells per well 16 h before infection. On the day of infection cultures were diluted 1:20. Once cultures had reached mid-log phase (OD600 0.6-0.8) at 2-3 h, 1 mL of bacteria were pelleted at 5,000 rpm for 3 min and washed 2X with PBS. Bacteria were diluted in serum-free DMEM (Hyclone) and added to cells at multiplicity of infection (MOI) of 5. Infected cells were spun at 1,000 rpm for 5 min then incubated for 10 min at 37 °C prior to adding fresh media. At indicated times post infection, cells were harvested with TRIzol for RNA isolation described below.

#### Vesicular stomatitis viral (VSV) infections

RAW 264.7 macrophages were seeded in 12-well tissue culture-treated plates at a density of 7×105 cells per well16 h before infection. The next day cells were infected with VSV-GFP virus at multiplicity of infection (MOI) of 1 in serum-free DMEM (HyClone). After 1 h of incubation with media containing virus, supernatant was removed, and fresh DMEM plus 10% FBS (Millipore) was added to each well. At indicated times post infection, cells were harvested with TRIzol for RNA isolation described below

#### *In vivo* Mtb infections

All infections were performed using procedures approved by Texas A&M University Institutional Care and Use Committee (AUP# 2021-0133). The Mtb inoculum was prepared as described above. Age- and sex-matched mice were infected via inhalation exposure using a Madison chamber (Glas-Col) calibrated to introduce 100-200 bacilli per mouse. For each infection, approximately 5 mice were euthanized immediately, and their lungs were homogenized and plated to verify an accurate inoculum. Infected mice were housed under BSL3 containment and monitored daily by lab members and veterinary staff. At the indicated time points, mice were euthanized, and tissue samples were collected. For cytokine transcript analysis, lungs were homogenized in 500 μL TRIzol, and RNA was isolated as described below.

#### Cell stimulation for immune activation and cell death

For immune activation RAW 264.7 macrophages were plated on 12-well tissue-culture treated plates at a density of 7.5×105 and allowed to rest overnight. Cells were then treated with lipopolysaccharide (LPS) from E. coli (Invivogen) at 100 ng/mL, or ISD (IDT, annealed in house) at 1 μg/mL, or IFNβ (PBL Assay Science) at 200 I/U per ml for the respective time points. Cells were collected for RNA isolation using TRIzol reagent. For IFNβ neutralization assays RAW 264.7 Srsf6 KD and SCR cells were treated for 24 h with IFNβ neutralizing antibody (PBL Assay Science) (1:250) prior to harvest with TRIzol. For cell death assays RAW 264.7 macrophages were plated on corning black 96-well half bottom plates at a density of 2.5×104 cells per well and allowed to rest overnight. Complete media was exchanged for complete media containing 5 μg/mL PI and 1 uM staurosporine (Tocaris Bioscience) or IFNβ neutralizing antibody (PBL Assay Science) (1:250). Total cell numbers were determined using NucBlue (Thermo Fisher) (2 drops per ml) in PBS with a subset of the plated cells. Nuclear incorporation of PI was measured by fluorescence at 4X magnification using a LionheartXF plate reader (BioTek) every 40 minutes for 24 h with 5% CO2 at 37 °C. For image analysis Gen 3.5 software (BioTek) was used.

#### mtDNA depletion

For 2′,3′-dideoxycytidine (ddC) depletion of mitochondrial DNA, RAW 264.7 macrophages were treated with 10 μM ddC and RNA was harvested after 8 days of culture with TRIzol.

#### Seahorse metabolic assays

Seahorse XF mito stress test kits and cartridges were prepared per Agilent’s protocols and analyzed on an Agilent Seahorse XF 96-well analyzer. The day before the assay *Srsf6* knockdown RAW 264.7 macrophages were seeded at 5×104 cells per well and rested overnight. Cells were processed per manufacturer’s directions and analyzed using the Agilent Seahorse Mito Stress Test kit (Agilent). Normalization was performed based on absorbance (Abs 562) of total protein concentration measured using a bicinchoninic acid assay (BCA) (Thermo Fisher Scientific). WAVE software was used for post-acquisition analysis.

#### RNA sequencing and analysis

RNA-seq analysis was performed on RAW 264.7 cells containing shRNA knockdowns of Srsf6, Srsf2, Srsf1, Srsf7, and Srsf9 compared to SCR control with biological triplicates of each cell line as described in (Wagner et al., 2021).

#### Alternative splicing analysis

Alternative splicing events were analyzed using Modeling Alternative Junction Inclusion Quantification (MAJIQ) and VOILA (a visualization package) with the default parameters described by Vaquero-Garcia et al., 2016. Uniquely mapped, junction-spanning reads were used by MAJIQ to construct splice graphs for transcripts by using the RefSeq annotation supplemented with *de-novo* detected junctions (de-novo refers to junctions that were not in the RefSeq transcriptome database but had sufficient evidence in the RNA-Seq data). The resulting gene splice graphs were analyzed for all identified local splice variations (LSVs). For every junction in each LSV, MAJIQ quantified expected percent spliced in (PSI) value in control and knockdown samples and expected change in PSI (ΔPSI) between control and knockdown samples. Results from VOILA were then filtered for high confidence changing LSVs (whereby one or more junctions had at least a 95% probability of expected ΔPSI of at least an absolute value of 10 PSI units between control and knockdown) and candidate changing LSVs (95% probability, 10% ΔPSI). For these high confidence results (ΔPSI 10%), the events were further categorized as single exon cassette, multi-exon cassette, alternative 5′ and/or 3′ splice site, or intron-retention.

#### ELISA

siRNA knockdown RAW 264.7 macrophages were plated in 12-well tissue culture-treated plates at 7.5×105 cells per well and rested overnight. The next day, plates were treated with 10 ng/mL LPS for 3 h (Invivogen) followed by 1 µg/mL poly dA:dT (Invivogen) or only the 10 ng/mL LPS for 4 h, and supernatants and RNA was collected using TRIzol. Cytokine levels of IL-1β, were determined using DuoSet ELISA Development Systems (R&D Systems) per manufacturer’s protocol using the undiluted cell culture supernatants.

#### Flow cytometry

For cell death and apoptosis assays, RAW 264.7 macrophages were plated in 12-well tissue culture-treated plates at 7.5×10^5^ cells per well and rested overnight. The next day cells were stimulated with 10 µM etoposide (Fisher), or 1µM ABT737 (ChemCruz), for the indicated time points prior to being lifted off culture plates with 1X PBS EDTA (Lonza). Single cell suspensions were made in 1X annexin binding buffer. Cells were stained for 5 min at RT in 5 µg/ml PI (Invitrogen), and 25 nM annexin-V (APC, eBioscience) and were then immediately analyzed on an LSR Fortessa X20 (BD Biosciences). PI fluorescence was measured under PE (585/15). For TMRE assays to assess mitochondrial membrane potential, cells were lifted from culture plates with 1X PBS EDTA (Lonza). Single cell suspensions were made in PBS 4% FBS (Millipore). Cells were stained for 20 min at 37 °C in 25 nM TMRE (Invitrogen), washed 1X in PBS 4% FBS (Millipore) and analyzed on an LSR Fortessa X20 (BD Biosciences). Flow-Jo software was used for post-acquisition analysis.

#### Cytoplasmic DNA enrichment

7×10^6^ RAW 264.7 macrophages were plated in 15 cm tissue culture-treated dishes and incubated at 37°C with 5%CO_2_. The next day cells were lifted with 1X PBS EDTA (Lonza) and resuspended in 5 mL PBS. Total DNA was isolated from 2% of resuspended cells, treated with 25 mM NaOH, boiled for 30 min, and then neutralized with 50 mM TRIS pH 8.0. The remainder of the cell suspension was pelleted at 3,000 rcf for 5 min. Cell pellets were resuspended in 500 μL cytosolic lysis buffer (50 mM HEPES pH 7.4, 150 mM NaCl, 50 μg/mL digitonin, 10 mM EDTA) and incubated on ice for 15 min. Cells were spun down at 1000 rcf to pellet intact cells and nuclei that were then used for obtaining the membrane fraction. The supernatant was transferred to a fresh tube and spun down at 15000 rcf to remove additional organelle fragments and transferred to a fresh tube again. Cytosolic protein was obtained by transferring 10% of supernatant to a fresh tube with 6X sample buffer + DTT and boiled for 5 min. Cytosolic DNA was isolated from the remaining supernatant by mixing an equal volume of 25:24:1 phenol: chloroform: isoamyl alcohol, with vigorous shaking followed by centrifugation for 10 min at ∼21130 rcf (max speed). The aqueous phase was transferred to a fresh tube and DNA was precipitated by mixing with 300 mM sodium acetate, 10 mM MgCl_2_, 1 μL glycogen, and 3 volumes of 100% ethanol, and then incubated overnight at −80°C. The precipitated DNA was pelleted by centrifugation at max speed for 20 min at 4 °C. The pellet was washed with 1 mL of cold 70% ethanol and centrifuged for 5 min at max speed, and then the pellet was air dried for 15 min at RT. The DNA was resuspended with 200μL DNase free water. For the mitochondrial membrane fraction, the pellet of intact cells, previously collected, was resuspended in 500μL membrane lysis buffer (50 mM HEPES pH 7.4, 150 mM NaCl, 1% NP-40), vortexed, and then centrifuged for 3 min at 7,000 rcf. 50 μL of the cleared lysate was transferred to a fresh tube 6X sample buffer + DTT was added. Samples were boiled 5 min. Immunoblotting was used to check for contaminating mitochondrial proteins in the cytosolic fraction compared to the membrane fraction by probing for mitochondrial ATP5A1. RT-qPCR was performed using total DNA diluted 1:100 and cytosolic DNA diluted 1:2. *Tert* (nuclear DNA), *CytB* and *Dloop* (mtDNA) were measured. Total and cytosolic fractions were normalized to *Tert* to control for variation in cell numbers.

#### Extracellular IFN-β assay

Macrophage-secreted type I IFN-β levels were determined using a L929 cells stably expressing a luciferase reporter gene under the regulation of type I IFN signaling pathway (L929 ISRE cells). 5×10^4^ L929 ISRE cells were seeded in a clear 96-well flat-bottomed plate (Thermo Scientific) and incubated at 37 °C with 5% CO_2_ the previous day. Macrophage cell culture media was collected, diluted 1:5 in complete media and transferred to the L929 ISRE cells and incubated for 6 h. Cells were washed with PBS, lysed in 30 μL cell culture lysis buffer, and transferred to a white 96-well flat-bottomed plate (Costar). 30 μL of Luciferase Assay System substrate solution (Promega) was added to the plate and luminescence read immediately using a Cytation5 plate reader (Biotek).

#### UV Crosslinking immunoprecipitation

9×10^6^ RAW 264.7 macrophages were seeded in 15cm tissue culture-treated plates and rested overnight. The next day treated with 1 mg/mL doxycycline for 24 h. Cells were washed with 1XPBS (Thermo Fisher) and UV treated at 2000μJoules ×100 with a UVstratalinker1800. Cell pellets were lysed in NP-40 lysis buffer with EDTA free protease inhibitor (Thermo Scientific) for 15 min and sonicated 3X for 10 min 30sec on/off (Biorupter). Lysates were treated with 200ng/mL RNaseA (Invitrogen) and 1U/mL RQ1 Dnase (Promega) and then incubated on 3XFLAG beads (Sigma Aldrich) for 3 h rotating at 4°C. Bound FLAG was eluted 3X using 20μL 5XFLAG peptide (Sigma Aldrich) by vortexing continuously for 15 min. Protein samples were collected and separated as described below and RNA was isolated using ethanol precipitation. Briefly, samples were treated with 0.1%SDS and 0.5 mg/mL proteinase K (Invitrogen). Samples were spun down at 10,000 rcf for 5 min at 4°C, and the pellet was vortexed with equal parts RNase free water and TRIzol, spun down at 10,000 rcf for 20 min at 4°C, and then pellets were stored in 100% ethanol +3M sodium acetate+ 1μL glycogen at −80°C overnight. Samples were spun down at 10,000 rcf for 10 min at 4°C and then the pellets were washed with 70% ethanol, spun down again, and then pellets were air dried for 15 min and RNA was resuspended in RNase free water. qPCR downstream analysis was performed as described below. No RT control cDNA samples were used to check for DNA contamination.

#### Immunofluorescence Microscopy

Cells were plated on glass coverslips in 24-well plates. At the designated time points, cells were washed with PBS (Thermo Fisher) and then fixed in 4% paraformaldehyde for 10 min at 37 °C. Cells were washed with PBS 3X and then permeabilized with 0.2% Triton-×100 (Thermo Fisher). Coverslips were incubated in primary antibody diluted in PBS + 5% non-fat milk + 0.2% Triton X-100 (PBS-MT) for 2h at RT. Primary antibodies used in this study were TOM20 clone 2F8.1(Millipore Sigma, 1:100); ANTI-FLAG M2 (Sigma Aldrich, 1:500); β-ACTIN (Abcam, 1:500) and DAPI (1:10,000). Coverslips were then washed 3X in PBS and incubated in secondary antibody (Invitrogen) diluted in PBS-MT for 1 h in the dark. Coverslips were then washed twice in PBS and then twice in deionized water. Following washes coverslips were mounted onto glass slides using ProLong Diamond antifade mountant (Invitrogen). Images were acquired on an Olympus Fluoview FV3000 Confocal Laser Scanning Microscope.

#### Protein quantification by immunoblot

Cells were washed with PBS and lysed in 1X RIPA buffer with protease and phosphatase EDTA free inhibitors (Thermo Scientific), with the addition of 1 U/mL Benzonase (Millipore) to degrade genomic DNA. Proteins were separated by SDS-PAGE on AnykD mini-PROTEAN TGX precast gel (Biorad) and transferred to 0.45μm nitrocellulose membranes (GE Healthcare). Membranes were blocked for 1h at RT in LiCOR Odyssey blocking buffer. Blots were incubated overnight at 4 °C with the following primary antibodies: β-ACTIN (Abcam, 1:5000), β-TUBULIN (Abcam, 1:5000) p-IRF3(S396) (Cell Signaling, 1:1000); IRF3 (Bethyl, 1:1000); ANTI-FLAG M2 (Sigma Aldrich, 1:5000); NWSHPQFEK (Genscript, 1:5000); CGAS (Cell Signaling, 1:1000); SRP55 (Bethyl, 1:1000); VIPERIN (RSAD2) (EMD Millipore, 1:1000); ATP5A1 (Bethyl, 1:1000); CYTOCHROME-C (Abcam, 1:1000). Membranes were washed 3X for 5 min in PBS-Tween20 and incubated with appropriate secondary antibodies (LI-COR) for 1h at RT prior to imaging on a LiCOR Odyssey Fc Dual-Mode Imaging System.

#### RNA isolation and qPCR analysis

For transcript analysis, cells and tissue were harvested in TRIzol and RNA was isolated using Direct-zol RNA Miniprep kits (Zymo Research) with 1 h DNase treatment. cDNA was synthesized with iScript cDNA Synthesis Kit (Bio-Rad). CDNA was diluted to 1:20 for each sample. A pool of cDNA from each treated or infected sample was used to make a 1:10 standard curve with each standard sample diluted 1:5 to produce a linear curve. RT-qPCR was performed using Power-Up SYBR Green Master Mix (Thermo Fisher) using a Quant Studio Flex 6 (Applied Biosystems). Samples were run in triplicate wells in a 384-well plate. Averages of the raw values were normalized to average values for the same sample with the control gene, *Actb*. To analyze fold induction, the average of the treated sample was divided by the untreated control sample, which was set at 1.

#### Semiquantitative qPCR analysis

cDNA was synthesized by iScript cDNA Synthesis Kit (Bio-Rad) using an extended 3 h amplification. Q5 high fidelity 2X Master mix (New England Biolabs) was used for PCR amplification using targeted primers. Loading dye was added to PCR products and samples were run on 2% agarose gel containing ethidium bromide at 100 volts for 1 h. Gels were imaged on LiCOR Odyssey Fc Dual-Mode Imaging System and bands were quantified.

#### Quantitation and Statistical Analysis

Statistical analysis of data was performed using GraphPad Prism software. Two-tailed unpaired Student’s t tests were used for statistical analyses, and unless otherwise noted, all results are representative of at least three biological samples (mean +/- SEM (n = 3 per group)). For *in vivo* data, Mann-Whitney statistical test was used (n=5). Agarose gel images for semi-quantitative RT-PCR and immunoblots are representative of n>3.

